# Dysregulation of ADAM10 shedding activity in naked mole-rat fibroblasts is due to deficient phosphatidylserine externalisation

**DOI:** 10.1101/2022.06.09.495538

**Authors:** Paulina Urriola-Muñoz, Luke A. Pattison, Ewan St. John. Smith

## Abstract

The naked mole-rat (NMR, *Heterocephalus glaber*) is of significant interest to biogerontological research, rarely developing age-associated diseases, such as cancer. The transmembrane glycoprotein CD44 is upregulated in certain cancers and CD44 cleavage by a disintegrin and metalloproteinase 10 (ADAM10) regulates cellular migration. Here we provide evidence that mature ADAM10 is expressed in NMR primary skin fibroblasts (NPSF), and that ionomycin increases cell surface ADAM10 localization. However, we observed an absence of ADAM10 mediated CD44 cleavage, as well as shedding of exogenous and overexpressed betacellulin in NPSF, whereas in mouse primary skin fibroblasts (MPSF) ionomycin induced ADAM10-dependent cleavage of both CD44 and betacellulin. Overexpressing a hyperactive form of the Ca^2+^-dependent phospholipid scramblase ANO6 in NPSF increased phosphatidylserine (PS) externalization, which rescued the ADAM10 sheddase activity and promoted wound closure in NPSF in an ADAM10-dependent manner. These findings suggest that dysregulation of ADAM10 shedding activity is due to a deficient PS externalization in NMR.

## INTRODUCTION

The naked mole-rat (NMR, *Heterocephalus glaber*) is a eusocial mammal with an exceptionally long lifespan of 35+ years, compared with sized rodents (*1*). Moreover, NMRs rarely develop age-associated diseases, such as cancer (*1*, *2*), which is why NMRs are recognized as species of particular biogerontological interest (*3*, *4*). However, what determines many aspects of NMR healthy ageing is unknown.

NMR cells undergo oncogenic transformation (*5*), and it is currently unclear where their cancer resistance stems from although it has been show that they exhibit a slow somatic mutation rate (*6*) and an altered immune response to carcinogenic insult (*7*). CD44 is a widely expressed, type I non-kinase transmembrane receptor for various extracellular matrix components, including hyaluronan, laminin and fibronectin (*8*, *9*). CD44 plays a role in numerous physiological and pathological processes, such as cellular adhesion, angiogenesis, metastasis, migration, and invasion (*10*–*14*). In addition, CD44 has been associated with cancer progression, being one of the most consistent markers of cancer stem cells and is involved in their generation, maintenance, and survival (*15*–*18*).

The principal domains of CD44 are the extracellular domain or ectodomain, the transmembrane domain, and the intracellular/cytoplasmic domain (*19*). The CD44 ectodomain can be cleaved by metalloproteases (*20*–*22*), a process stimulated by extracellular Ca^2+^ influx, the activation of Rho family small GTPases, Rac and Ras oncoproteins, and the activation of protein kinase C (*22*–*24*). Cleavage of the CD44 ectodomain is highly prevalent in patients with tumours including glioma, breast, lung, colon and ovarian cancers (*25*, *26*), and high levels of soluble CD44 have been found in the serum of patients with cancer (*27*, *28*). Furthermore, CD44 cleavage promotes tumour cell migration (*24*, *29*–*31*). Therefore, it is clear that CD44 cleavage plays an important role in tumour progression.

A Disintegrin and Metalloprotease 10 (ADAM10) is one of the metalloproteases involved in CD44 cleavage (*32*, *33*). ADAM proteins are type I metalloproteases responsible for the shedding of different membrane-bound receptors and ligands controlling many cellular functions (*34*). ADAM10 is one of best characterized ADAM family members, cleaving many proteins, including CD44, betacellulin (BTC) and Notch (*22*, *35*, *36*), as well as being involved in diverse physiological processes, such as fertilization, neurogenesis and angiogenesis (*35*, *37*). ADAM10 dysregulation is associated with different pathologies, such as embryonic lethality in mice due to disturbed Notch signalling being caused by depletion of the ADAM10 gene (*36*) and upregulation of ADAM10 occurring in melanoma (*38*), as well as roles for dysregulation of ADAM10 shedding activity occurring in Alzheimer’s disease, and cancer development (*39*, *40*).

Constitutive ADAM10 shedding activity is enhanced by Ca^2+^ influx, which can be induced by ionophores as ionomycin (IM) (*41*, *42*), as well as exposure of the negatively charged phosphatidylserine (PS) in the outer leaflet of the cell membrane (*43*).

Considering the role of ADAM10 in cellular function, we hypothesize that another mechanism that might contribute to NMR cancer resistance would be dysregulation of ADAM10 shedding activity. In experiments conducted in NMR primary skin fibroblasts (NPSF), we report an absence of CD44 cleavage induced by IM. Furthermore, although IM induced the translocation of ADAM10 to the cell membrane of NPSF, it does not induce ADAM10-dependent shedding of the ADAM10 substrate BTC when overexpressed. However, when NPSF overexpressed a hyperactive form of anoctamin 6 (ANO6-HA, a Ca^2+^-dependent phospholipid scramblase), an increase in PS externalization was observed alongside an IM induced, ADAM10-dependent BTC shedding. Finally, ANO6-HA overexpression promoted ADAM10-dependent wound closure. All these data suggest that a deficiency in PS externalization may contribute to a dysregulation of ADAM10 shedding activity in NPSF.

## MATERIALS AND METHODS

### Animals

Experiments were performed on cells isolated from a mixture of male and female C57BL6/J mice (8-15 weeks old), and a mixture of male and female, non-breeder NMRs (7-43 months old). Mice were conventionally housed with nesting material and a red plastic shelter in temperature-controlled rooms at 21° C, with a 12 h light/dark cycle and access to food and water ad libitum. Naked mole-rats were bred in-house and maintained in an inter-connected network of cages in a humidified (~55 %) temperature-controlled room at ~28° C, with red lighting (08:00-16:00) and had access to food ad libitum. In addition, a heat cable provided extra warmth under 2-3 cages/colony. Experiments were conducted under the Animals (Scientific Procedures) Act 1986 Amendment Regulations 2012 under a Project License (P7EBFC1B1) granted to E. St. J. Smith by the Home Office and approved by the University of Cambridge Animal Welfare Ethical Review Body.

### NMR and mouse primary skin fibroblast isolation

NMR and mouse primary skin fibroblast (NPSF and MPSF) isolation were performed as described before (*5*). Briefly, NMR were humanely killed by CO_2_ exposure followed by decapitation, whereas mice were killed by cervical dislocation of the neck and cessation of circulation. Skin tissue was collected from each animal on ice-cold PBS 1X (#70011-044; Gibco). For NPSF, skin was taken from the underarm, dorsal and ventral surfaces of each animal, and was cleaned of any fat or muscle tissue and sprayed with 70% ethanol. For MPSF, both external ears were collected from each animal and was cleaned of any hair and sprayed with 70% ethanol. The mouse pinna lacks significant hair, like NMR skin in general, but NMR do not have pinnae and hence it was not feasible to isolate fibroblasts from glabrous skin of identical origins in both species; NPSF grow slowly and thus isolating from multiple regions enabled experiments to be conducted in a timely manner, as well as minimising the number of animals used, and when tracking the area from where fibroblasts were isolated no difference between experiments was found. Once isolated and cleaned, tissue was washed twice with PBS 1X and finely minced with sterile blades (#BS2982; Swann Morton). For dissociation, minced skin was incubated at 37°C for 3 – 5 hours in 5 ml of DMEM high glucose (#41965-039; Gibco) supplemented with 10 mg ml^-1^ collagenase (#C9891; Sigma-Aldrich), 1000 units ml^-1^ hyaluronidase (#H3506; Sigma-Aldrich). The tissue was briefly vortexed every 30 minutes. Cells were subsequently pelleted by centrifuging at 500 g for 3 minutes and resuspended in PBS 1X, then the cells were centrifuged again and resuspended in cell culture medium: DMEM high glucose (#41965-039; Gibco) supplemented with 15% fetal bovine serum (#F7524-500ML; Sigma-Aldrich), 100 units ml^-1^ penicillin, 100 μg ml^-1^ streptomycin (#15140122; Gibco) and 100 μg ml^-1^ Primocin (#ant-pm-2; InvivoGen). NMR culture media was further supplemented with 1X non-essential amino acids (#11140-050; Gibco) and 1 mM sodium pyruvate (#11360-039; Gibco). This cell suspension was passed through a 70 μm filter (#352350; Falcon) and seeded on a treated cell culture flask (T-75 #658175; Greiner Bio-One). NPSF cultures were incubated in a humidified 32 °C incubator with 5% CO_2_ and 3% O_2_, whereas MPSF cultures were incubated in a humidified 37 °C incubator with 5% CO_2_.

### Mouse and NMR SV40-Ras skin fibroblasts

Transformed mouse and NMR SV40-Ras skin fibroblasts that were previously developed in our lab (*5*) were used. NMR and mouse SV40-Ras skin fibroblasts were cultured in DMEM high glucose (#41965-039; Gibco) supplemented with 15% fetal bovine serum (#F7524-500ML; Sigma-Aldrich) and 100 units ml-1 penicillin, 100 μg ml-1 streptomycin (#15140122; Gibco). NMR culture media was further supplemented with 1X non-essential amino acids (#11140-050; Gibco) and 1 mM sodium pyruvate (#11360-039; Gibco). Mouse cells were incubated in a humidified 37 °C incubator with 5% CO_2_. NMR cells were incubated in a humidified 32 °C incubator with 5%CO_2_ and 3% O_2_.

### Protein Extraction and Western Blotting

Protein extraction was performed by homogenizing cells to produce a cell lysate in a buffer containing 150 mM NaCl, 10 mg ml^-1^ PMSF, 60 mM DTT, 1% NP-40, 1 mM sodium orthovanadate and 50 mM Tris–HCl pH 7.4, plus a general metalloprotease inhibitor BB-94 10 μM (#196440; Calbiochem) and protease inhibitor cocktail, which included AEBSF (2 mM), aprotinin (0.3 μM), bestatin (116 μM), E-64 (14 μM), leupeptin (1 μM) and EDTA (1 mM) (#P2714; Sigma-Aldrich), and then centrifuging for 10 min at 16,100 x g at 4°C. The samples were run on a 15% polyacrylamide gel (SDS–PAGE) under reducing and denaturing conditions, and then transferred to nitrocellulose membrane (#88018; Thermo Fisher). The nitrocellulose membrane was blocked with 3% (w/v) non-fat milk (#70166; Sigma-Aldrich), 0.1% Tween-20 in TBS, pH 7.4 and then incubated overnight at 4°C with one of the following antibodies: anti-CD44 (0.04 μg ml^-1^, #ab157107; Abcam), generated against the C-terminal and able to detect different shedding products of CD44 between 15 and 25 kDa (*29*, *44*), anti-ADAM10 (0.04 μg ml^-1^, #PA5-87899; Invitrogen), anti-ANO6 (0.5 μg ml^-1^, #PA5-35240; Invitrogen) or, as a loading control, anti-β-actin (8H10D10) (1:10,000, cat#3700S; Cell Signaling). Membranes were then incubated with a donkey anti-rabbit IgG secondary antibody conjugated with IRDye^®^ 800CW (0.2 μg ml^-1^, #926-32213; LI-COR) and donkey anti-mouse IgG secondary antibody conjugated with IRDye^®^ 680RD (0.2 μg ml^-1^, #926-68072; LI-COR) in blocking solution for 1 h at room temperature. Protein bands were evaluated using the Oddysey^®^XF Imaging System (LI-COR) and images were analyzed using the Image Studio Lite software (LI-COR).

### Membrane protein biotinylation

Fibroblasts were grown in a 10 cm dish (#CC7682-3394; Cyto-one). MPSF and NPSF were treated for 1 h with IM 0.5 μM or DMSO as vehicle in DMEM high glucose (#41965-039; Gibco). Mouse and NMR SV40-Ras skin fibroblasts were treated for 15 and 10 min with IM 2.5 μM, respectively, or DMSO as vehicle in DMEM high glucose (#41965-039; Gibco). After treatment, cells were washed twice with Hanks-Ca^2+^ medium pH 7.4. Cells were then incubated with EZ-Link™ Sulfo-NHS-SS-Biotin (0.5 mg ml^-1^, #21331; Thermo Fisher) for 30 min at 4°C. Biotin was washed three times with Hanks-Ca^2+^ supplemented with 1 mg ml^-1^ glycine. Subsequently, cells were recollected in a solution of distilled water containing a protease inhibitor cocktail as described above (#P2714; Sigma-Aldrich) and centrifuged at 16,100 x g for 2 min at 4°C. The pellet was resuspended in a buffer containing 150 mM NaCl, 10 mg/ml PMSF, 60 mM DTT, 1% NP-40, 1 mM sodium orthovanadate and 50 mM Tris–HCl pH 7.4, plus a general metalloprotease inhibitor BB-94 10 μM (#196440; Calbiochem) and protease inhibitor cocktail as described above (#P2714; Sigma-Aldrich). The protein concentration (total fraction) was calculated, and 200 μg of protein was precipitated with NeutrAvidin™ Agarose Resins (#29201; Thermo Fisher Scientific) after an incubation of 1 h at 4°C with occasional manual agitation. The pellet was washed with Hanks without Ca^2+^ supplemented with 0.1% SDS and 1% NP-40 and then centrifuged at 16,100 x g for 2 min at 4°C and resuspended with 20 μl Hanks-Ca^2+^ supplemented with 1 mg ml^-1^ glycine, pH 2.8. Then the sample was centrifuged in the same conditions above and the supernatant was rescued (this was the biotinylated fraction). All fractions were run on a 10% polyacrylamide gel (SDS– PAGE) under reducing and denaturing conditions.

### Immunofluorescence

MPSF and NPSF were grown in a MatTek (#P35GC-1.5-14-C; MatTek life sciences) dish under the conditions explained above. The localization of ADAM10 was assayed in fibroblasts treated for 1 h with IM 0.5 μM or DMSO as vehicle. Fibroblasts were fixed in 100% methanol for 15 min at −20°C and then permeabilized with 0.1% Triton X-100. Samples were blocked with 3% BSA (#A2153; Sigma-Aldrich) in TBS containing 0.1% Tween-20 (T-TBS) for 1 h at room temperature. A primary antibody against ADAM10 (#PA5-87899; Invitrogen, host - rabbit) was applied at 1 μg ml^-1^ diluted in bocking solution and dishes were incubated overnight at 4°C in a humidified chamber. The next day, the dish was washed three times for 10 min in T-TBS. Goat anti-rabbit IgG (H+L) Cross-Adsorbed Secondary Antibody, Alexa Fluor™ 488 (#A-11008; Invitrogen) at 1 μg ml^-1^ diluted in 3% BSA-T-TBS was added to the dish and then incubated for 1 h at room temperature, before washing dishes as before. Subsequently, 5 μg ml^-1^ of wheat germ agglutinin was added and incubated for 10 min at 37°C. Then the dishes were washed as before and NucRed™ Live 647 (#R37106; Invitrogen) was added to the dish and incubated for 5 min at room temperature. Finally, dishes were washed as above and cells observed using a Leica SP5 laser-scanning confocal microscope with 63x oil objective. Images were acquired in sequential scan mode: wheat germ agglutinin fluorescence was excited with a 405 nm laser and emission 415 - 475 nm recorded, Alexa Fluor™ 488 was excited with a 488 nm laser and emission between 498 - 590 nm captured, a 633 nm laser was used to excite NucRed™ Live 647 and emission between 645 - 633 nm detected. Composite images were created in Image J.

### Cell transfection

Mouse and NMR primary and SV40-Ras skin fibroblasts were transfected with plasmids encoding (AP)-tagged BTC (*45*) or with the hyperactive ANO6 (ANO6-HA) (*43*). Briefly, fibroblasts were seeded to 80% confluence and the following day were washed with PBS 1X and cultured in DMEM high glucose (#41965-039; Gibco) using the same conditions described above. Cells were cultured for 24 h with a complex of DNA and FuGENE^®^ HD Transfection Reagent (#E2311; Promega) at a ratio of 1:3 prepared in Opti-MEM reduced serum medium (#31985070; Gibco).

### ADAM10 shedding assay

For the shedding assay, fibroblasts were seeded in 12 well plates (#3513; Corning Costar). Every condition of the experiment was performed in non-transfected and transfected cells with the (AP)-tagged BTC plasmid. After 24 h, cells were washed once in PBS 1X and incubated for 30 min in Opti-MEM reduced serum medium (#31985070; Gibco). Then, treatment with IM and inhibitors was performed as indicated in the results section. After the treatment, conditioned media were collected and cells lysed with a solution containing 2.5% Triton X100 (#T8787; Sigma-Aldrich), 1 mM 1-10 phenanthroline (#131377; Sigma-Aldrich) and 1 mM EDTA (#10618973; Fisher Scientific). 100 μl of conditional media and 10 μl of cell lysate were added into 96 well plates (#E2996-1610; Starlab). Then 90 μl of AP-buffer (100 mM Tris, 100 mM NaCl, 20 mM MgCl_2_, pH 9.5) was added only to the cell lysate wells. After that, both conditional media and cell lysate, were incubated with 100 μl of 2 mg ml^-1^ p-NPP (p-nitrophenyl phosphate) substrate (#34045; Thermo Fisher) diluted in AP-buffer. Then the plate was incubated for 1 h at 37°C. The AP activity was measured by a spectrophotometer (CLARIOstar, BMG Labtech) at 405 nm. Three identical wells were prepared, and the ratio of AP activity in the medium and that of the cell lysate plus medium was calculated. For stimulated shedding or vehicle, the fold increase in the ratio of AP activity obtained after stimulation is shown relative to ratio of AP activity in control wells. Each experiment was conducted at least three times. Transfection efficiency was controlled by determining the ratio of the activity of AP in the medium over the AP-activity in the cells plus the medium.

### PS quantification in the outer leaflet of the cell membrane

20,000 fibroblasts per well were seeded in a white 96 well plate (#CC7682-7596; CytoOne). The following day, cells were washed with PBS 1X and incubated with Imaging media (140 mM NaCl, 2.5 mM KCl, 1.8 mM CaCl_2_, 1 mM MgCl_2_, 20 mM HEPES) supplemented with 15% of fetal bovine serum (#F7524-500ML; Sigma-Aldrich). PS exposure was evaluated through the ratio between the Annexin V and number of cells. Annexin V binding to PS was measured using the RealTime-Glo™ Annexin V luminescence assay (#JA1000; Promega), following the manufacturer’s instructions, binding of PS to annexin V generating a luminescence signal. The relative luminescence unit (RLU) was measured using a luminescence microplate reader (CLARIOstar, BMG Labtech). To determine the number of cells, after measurement of the RLU, cells were washed once with PBS 1X and then incubated with NucRed™ Live 647 ReadyProbes^®^ reagent (Invitrogen #R37106) in Imaging media for 15 min at room temperature. Cells were then washed with PBS 1X and the relative fluorescence units (RFU) was measured using a fluorescence microplate reader (CLARIOstar, BMG Labtech). Three identical wells were prepared, and the ratio of RLU/RFU was calculated.

### Wound closure assay

20,000 fibroblasts were plated in each well of a culture-insert 2 well in μ-dish 35 mm, high (#81176; Ibidi) and transfected as mentioned above. The following day, the silicon insert was removed from the dish and cells were washed once with PBS 1X and DMEM high glucose media (#41965-039; Gibco), supplemented with 5% of fetal bovine serum (#F7524-500ML; Sigma-Aldrich). During all the experiments, fibroblasts were culture in the conditions mentioned above. To measure the wound closure percentage, 4 pictures from each dish were taken at 0, 3, 6, 9, 12, 24, 36 and 48 h using a 5MP USB 2.0 Color CMOS C-Mount Microscope Camera (MU500-CK-3PL; AmScope) and the wound closure area was calculated using the TScratch software (*46*).

## RESULTS

### CD44 cleavage is not induced by IM in NMR primary skin fibroblasts

CD44 cleavage by metalloproteases has been implicated in cancer cell migration (*21*, *22*, *29*, *47*) and cancer resistance is a characteristic of naked mole-rat (NMR) biology (*2*). We therefore used immunoblotting to analyse the extent of IM-induced CD44 cleavage by ADAM10 in NMR primary skin fibroblasts (NPSF) and mouse primary skin fibroblasts (MPSF); IM is a well characterised initiator of Ca^2+^ influx that activates ADAM10 sheddase activity (*41*).

In all conditions, cells were preincubated for 30 min with the proteasomal inhibitor MG-132 (10 μM) to prevent degradation of CD44 cleavage products (*48*). In MPSF, the CD44 full length protein had a molecular weight of ~100 kDa, comparable to that previously published (*29*, *47*). After 1 h treatment with 0.5 and 1 μM IM, increased density of two fragments between ~15 and ~25 kDa was observed in MPSF (Fig. S1A). To determine the participation of metalloproteases, and specifically ADAM10 in this process, the effects of the following drugs were observed: batimastat, a general metalloprotease inhibitor (BB-94, 10 μM), GW280264X, an inhibitor of ADAM10/ADAM17 (GW, 1 μM) and GI254023X, which is selective for ADAM10 (GI, 1μM) (*49*). In MPSF, all three inhibitors prevented IM-induced CD44 cleavage (Fig. 1A), suggesting a key role for ADAM10 in CD44 cleavage in MPSF. By contrast, in NPSF, IM failed to induce CD44 cleavage using the same IM concentrations and incubation times that IM induces CD44 cleavage in MPSF (Fig. 1B and S1B). These data show that CD44 cleavage is not induced by IM in NPSF.

**Fig. 1.**
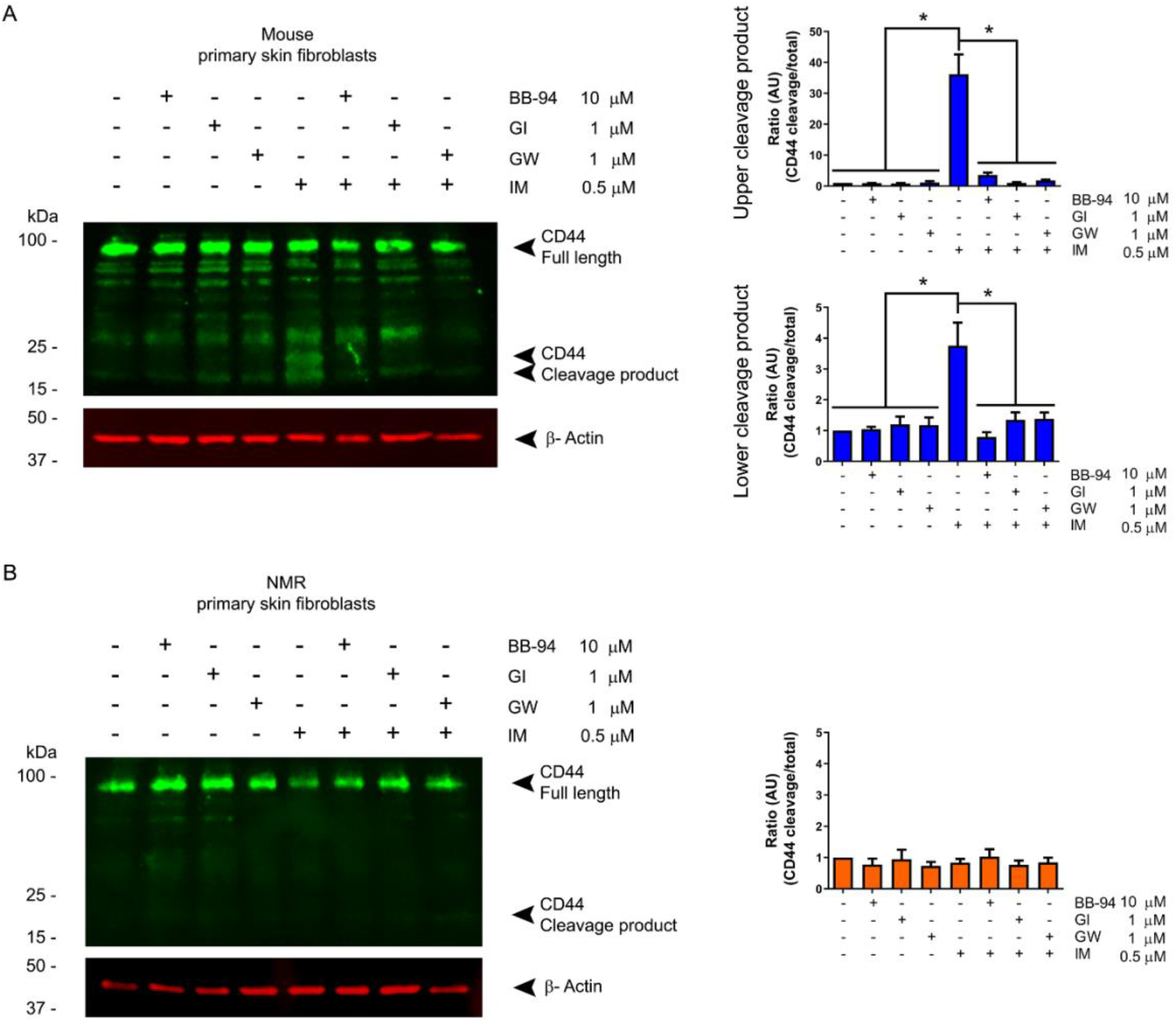
CD44 cleavage is not induced by IM in NPSF. An immunoblot was performed to evaluate the full length and cleaved products of CD44 in **(A)** MPSF and **(B)** NPSF. Cells were treated for 30 min with a general inhibitor of metalloproteases (batimastat, BB, 10 μM), an ADAM10 inhibitor (GI254023X, GI, 1 μM) or an ADAM10/ADAM17 inhibitor (GW280264X, GW, 1 μM), before stimulating cells for 1 h with ionomycin (IM, 0.5 μM). (A) In MPSF, the full CD44 protein has a molecular weight of ~100 kDa, when CD44 is cleaved, two fragments between ~15 and ~25 kDa are produced. Comparison of the upper and lower cleavage products with the full protein level generated a ratio of CD44 cleavage/total. (B) In NPSF, the full CD44 protein has a molecular weight of ~100 kDa, when CD44 is cleaved, only one fragment of ~15 kDa was observed. Comparison of the cleavage product with the full protein level generated a ratio of CD44 cleavage/total. Vehicle: DMSO. Results are mean ± SEM; n=3. For statistical analysis mean values were compared using an ANOVA and Tukey’s post-hoc test; *p ≤ 0.05.

**Fig. S1.**
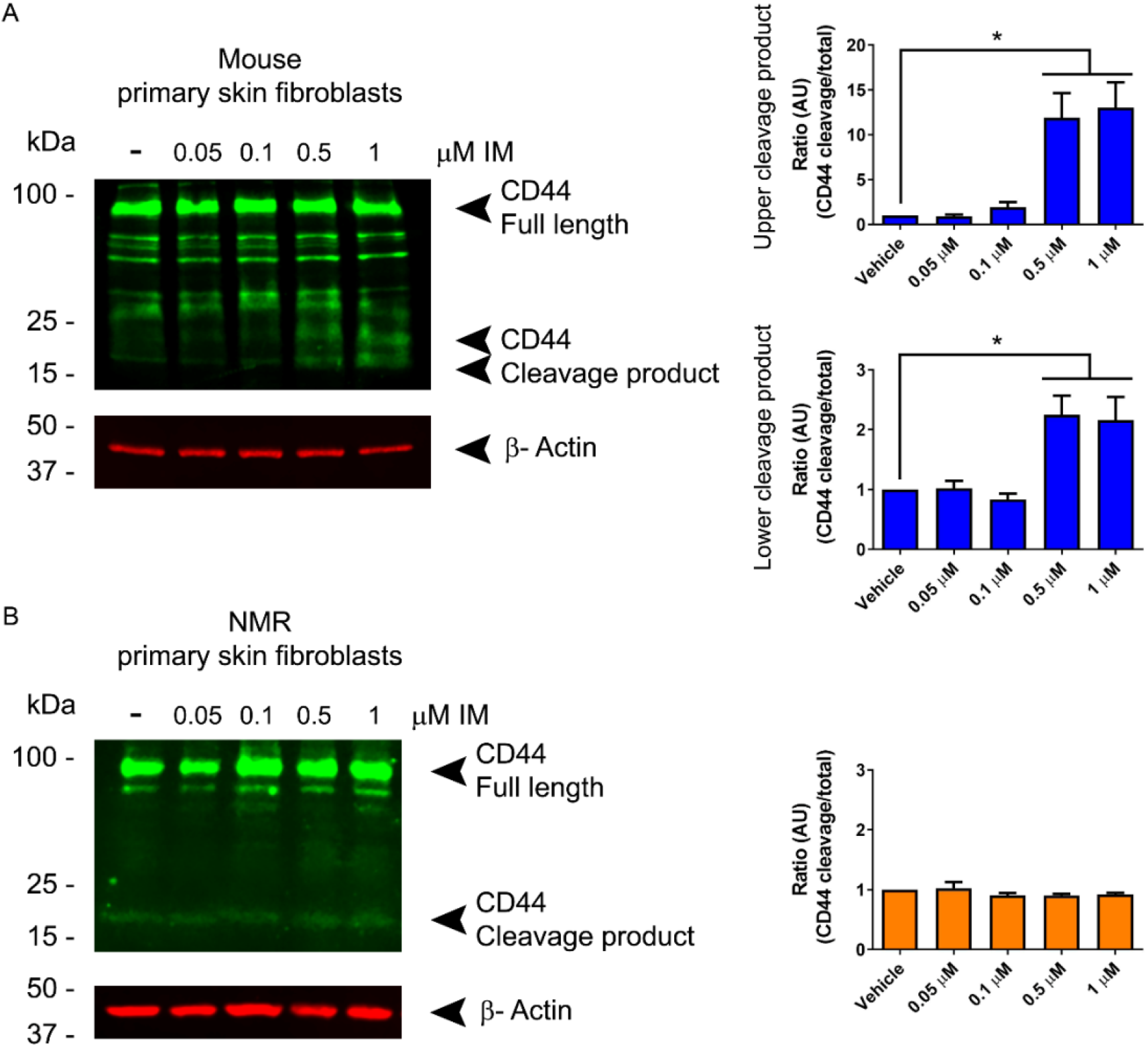
CD44 cleavage is induced by IM in MPSF but not in NPSF. An immunoblot was performed to evaluate the full length and cleaved products of CD44 in **(A)** MPSF and **(B)** NPSF. Cells were treated for 1 h with 0.05, 0.1, 0.5 and μM IM. **(A)** In MPSF, the full CD44 protein has a molecular weight of ~100 kDa, when CD44 is cleaved, two fragments between ~15 and ~25 kDa are produced. Comparison of the upper and lower cleavage products with the full protein levels generated a ratio of CD44 cleavage/total. **(B)** In NPSF, the full CD44 protein has a molecular weight of ~100 kDa, when CD44 is cleaved, only one fragment of ~15 kDa was observed. Comparison of the cleavage product with the full protein levels generated a ratio of CD44 cleavage/total. Vehicle: DMSO. Results are mean ± SEM; n=3. For statistical analysis mean values were compared using an ANOVA and Tukey’s post-hoc test; *p ≤ 0.05.

### NPSF express ADAM10, which undergoes cell membrane translocation, but is not activated by IM

To determine if the lack of IM-induced CD44 cleavage in NPSF is due to lack of ADAM10 expression, we compared ADAM10 expression levels in NPSF and MPSF. Pro- and mature ADAM10 were observed in both MPSF and NPSF, higher levels of pro-ADAM10, but not mature ADAM10, being expressed in NPSF compared to MPSF (Fig. 2A). Because CD44 cleavage takes place at the cell surface, we next evaluated if IM induced ADAM10 cell membrane translocation. MPSF and NPSF were treated for 1 h with 0.5 μM IM and the surface localisation of ADAM10 was evaluated by biotinylation. An increase in mature ADAM10 was found in both MPSF and NPSF treated with IM compared with the vehicle in the biotinylated fraction, with no change in the levels of mature ADAM10 protein in the total fraction (Fig. 2B). These results were confirmed with immunostaining of ADAM10 in MPSF and NPSF, where co-localisation of ADAM10 and wheat germ agglutin labelling of the cell membrane was conducted (Fig. 2C). Finally, to evaluate ADAM10 shedding activity more broadly, MPSF and NPSF were transfected with the alkaline phosphatase (AP)-tagged ADAM10 substrate betacellulin (BTC) (*35*). A significant increase in BTC shedding after 1 h of incubation with IM 0.5 μM occurred in MPSF, which was prevented with batimastat (BB-94, 10 μM), GI254023X (GI, 1 μM), or GW280264X (GW, 1 μM) (Fig. 2D), suggesting that ADAM10 participates in the IM-induced BTC shedding in MPSF. However, IM did not induce BTC shedding in NPSF (Fig. 2D). Therefore, using two different approaches, evaluating the cleavage of an endogenous substrate, CD44 (Fig. 1B) or the shedding of an exogenous and well-studied substrate, BTC (Fig. 2D), IM failed to induce ADAM10-dependent shedding activity in NPSF.

**Fig. 2.**
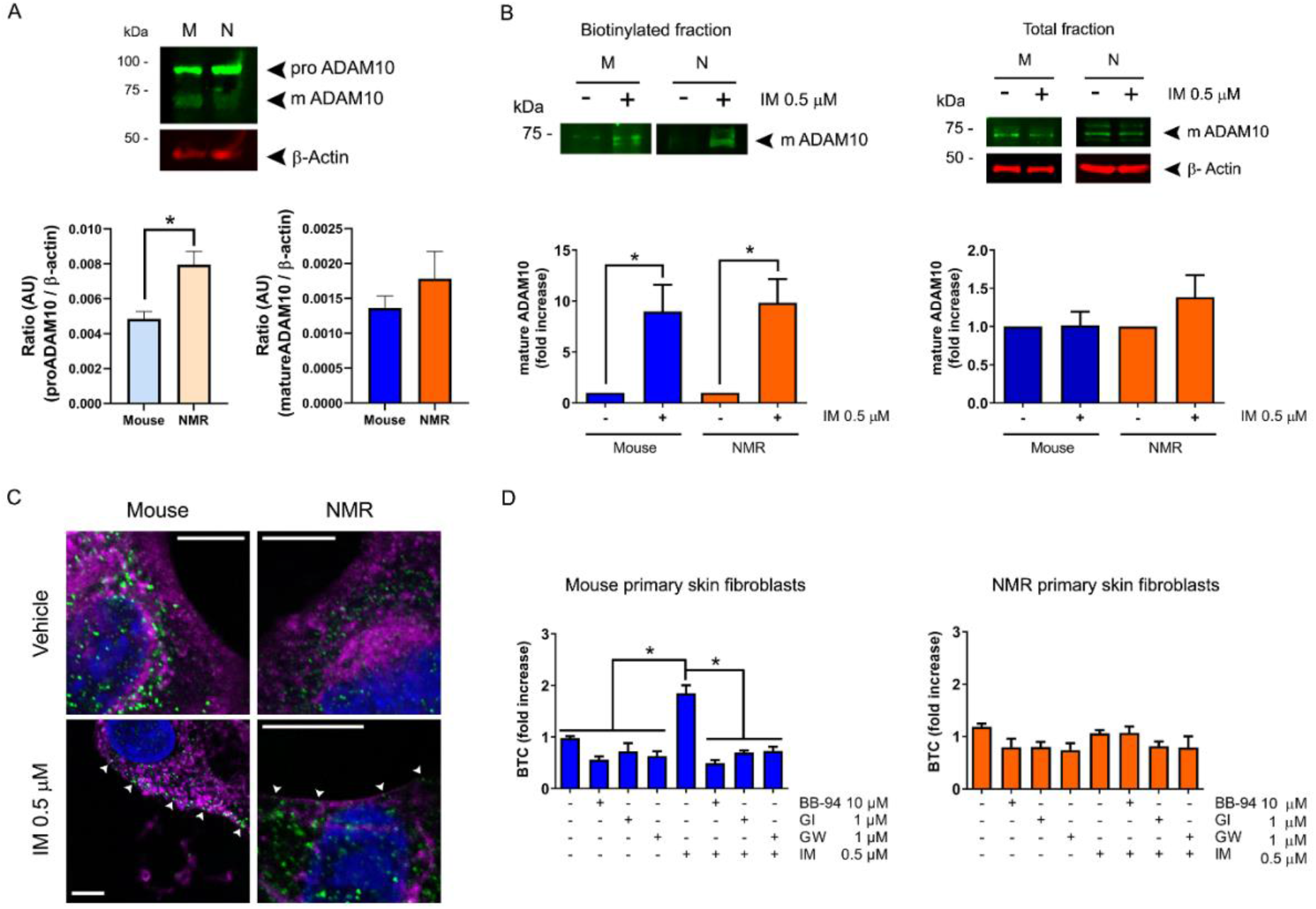
NPSF express ADAM10, which undergoes cell membrane translocation, but is not activated by IM. **(A)** Immunoblot of ADAM10, where pro-ADAM10 (pro) and mature ADAM10 (m) were observed in both MPSF (M) and NPSF (N). β-actin was used as loading control. Results are mean ± SEM; n=3. For statistical analysis, mean values were compared using an unpaired t test; *p ≤ 0.05. **(B)** Immunoblot of cell surface and total mature ADAM10 protein fraction in MPSF and NPSF treated for 1 h with ionomycin (IM, 0.5 μM). β-actin was used as loading control in the total protein fraction. Results are mean ± SEM; n=3. For statistical analysis, mean values were compared using an unpaired t test; *p ≤ 0.05. **(C)** Pseudocoluor; ADAM10 (green), wheat germ agglutinin (magenta) and NucRed™ Live 647 (blue) staining in MPSF and NPSF, with or without a treatment for 1 h with ionomycin (IM, 0.5 μM). White arrowheads indicate green puncta of ADAM10 at the cell membrane following IM treatment. Vehicle: DMSO. Scale bar: 10 μm. **(D)** MPSF and NPSF were transfected with the AP-tagged ADAM10 substrate BTC. BTC shedding was evaluated after cells were treated for 30 min with a general inhibitor of metalloproteases (batimastat, BB, 10 μM), an ADAM10 inhibitor (GI254023X, GI, 1 μM) or an ADAM10/ADAM17 inhibitor (GW280264X, GW, 1 μM), and then cells were stimulated for 1 h with ionomycin (IM, 0.5 μM). Vehicle: DMSO. Results are mean ± SEM; n=3. For statistical analysis, mean values were compared using an ANOVA and Tukey’s post-hoc test; *p ≤ 0.05.

### IM induces CD44 cleavage and ADAM10 shedding activity in both mouse and NMR SV40-Ras skin fibroblasts

To determine if the absence of IM-induced ADAM10 shedding activity is a general phenomenon of NMR cells, or is restricted to primary cells, we conducted experiments using transformed mouse and NMR SV40-Ras skin fibroblasts previously developed in our lab (*5*). Firstly, IM-induced CD44 cleaved protein levels were determined by immunoblot. In both mouse and NMR SV40-Ras skin fibroblasts, 0.5 μM IM for 1 h did not induce an increase of CD44 cleavage products (data not shown). Therefore, we evaluated if CD44 cleavage could be induced by 2.5 μM IM, a concentration previously reported to induce ADAM10 shedding activity in mouse embryonic fibroblasts (*50*), at 5, 10, 15, and 30 min. A significant increase of two fragments between ~15 and ~25 kDa was observed after 15 min treatment with 2.5 μM IM in mouse cells (Fig. S2A) and after 10 min treatment in NMR cells (Fig. S2B). In SV40-Ras skin fibroblasts of both species, CD44 cleavage was prevented by batimastat (BB-94, 10 μM), GI254023X (GI, 1 μM) or GW280264X (GW, 1 μM) (Fig. 3, A and B), suggesting that ADAM10 participates in IM-induced CD44 cleavage in both mouse and NMR SV40-Ras cells.

**Fig. 3.**
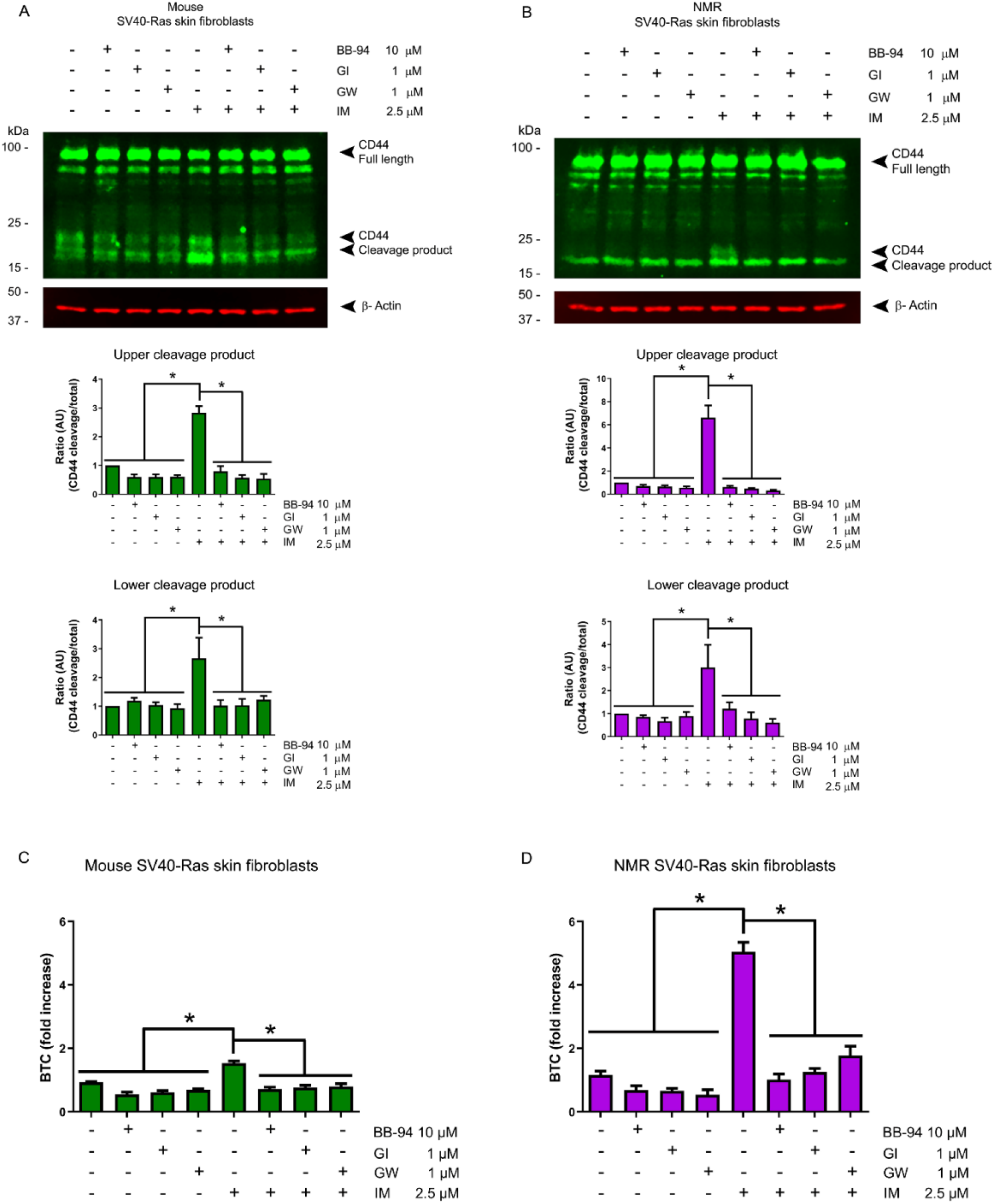
IM induces CD44 cleavage and ADAM10 shedding activity in both mouse and NMR SV40-Ras skin fibroblasts. Immunoblot evaluation of full length and cleavage products of CD44 in **(A)** mouse and **(B)** NMR SV40-Ras skin fibroblasts. Cells were treated for 30 min with a general inhibitor of metalloproteases (batimastat, BB, 10 μM), an ADAM10 inhibitor (GI254023X, GI, 1 μM) or an ADAM10/ADAM17 inhibitor (GW280264X, GW, 1 μM), before stimulating cells for with ionomycin (IM, 2.5 μM). Full CD44 protein has a molecular weight of ~100 kDa, when CD44 is cleaved, two fragments between ~15 and ~25 kDa are produced. The upper and lower cleavage product is compared with the full protein levels to generate a ratio of CD44 cleavage/total. β-actin was used as loading control. Vehicle: DMSO. Results are mean ± SEM; n=3. For statistical analysis, mean values were compared using an ANOVA and Tukey’s post-hoc test; *p ≤ 0.05. **(C)** Mouse and **(D)** NMR SV40-Ras skin fibroblasts were transfected with the AP-tagged ADAM10 substrate BTC. BTC shedding was evaluated after cells were treated for 30 min with a general inhibitor of metalloproteases (batimastat, BB, 10 μM), an ADAM10 inhibitor (GI254023X, GI, 1μM) or an ADAM10/ADAM17 inhibitor (GW280264X, GW, 1 μM), and then cells were stimulated with IM, 2.5 μM. Vehicle: DMSO. Results are mean ± SEM; n=3. For statistical analysis, mean values were compared using an ANOVA and Tukey’s post-hoc test; *p ≤ 0.05.

In addition, pro- and mature ADAM10 is expressed in both, mouse and NMR SV40-Ras skin fibroblasts, with higher pro-ADAM10, but not mature ADAM10, levels in mouse compared to NMR cells (Fig. S3A). An IM-induced increase in mature ADAM10 was found in the biotinylated cell membrane protein fraction of both mouse and NMR SV40-Ras skin fibroblasts, with no change detected in the total fraction (Fig. S3B). Similar to results obtained when measuring IM-induced CD44 cleavage, a significant increase in the shedding of BTC after incubation with 2.5 μM IM was observed in both mouse and NMR SV40-Ras skin fibroblasts transfected with the AP-tagged ADAM10 substrate BTC. This increase was prevented with the batimastat (BB-94, 10 μM), GI254023X (GI, 1 μM), or GW280264X (GW, 1 μM) (Fig. 3, C and D). Overall, these data suggest that IM activated the shedding activity of ADAM10, inducing ADAM10-dependent CD44 cleavage and BTC shedding in transformed cells from both mouse and NMR, thus demonstrating that NMR ADAM10 is functional in certain conditions.

**Fig. S2.**
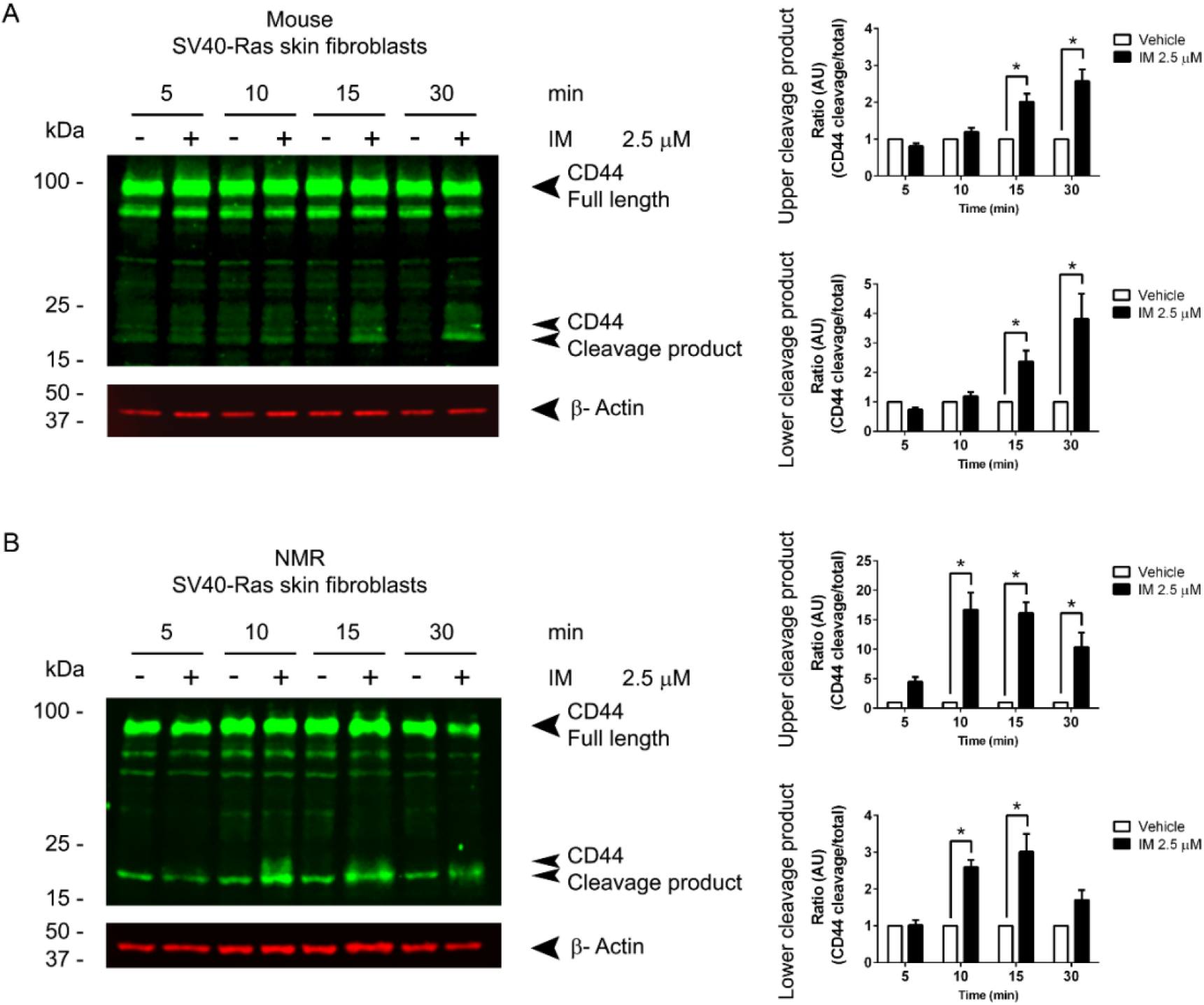
CD44 cleavage is induced by IM in transformed mouse and NMR SV40-Ras skin fibroblasts. An immunoblot was performed to evaluate the full length and cleaved products of CD44 in transformed **(A)** mouse and **(B)** NMR SV40-Ras cells. Cells were treated for 5, 10, 15 or 30 min with 2.5 μM IM. In **(A)** mouse and **(B)** NMR SV40-Ras cells, the full CD44 protein has a molecular weight of ~100 kDa, when CD44 is cleaved, two fragments between ~15 and ~25 kDa are produced. Comparison of the upper and lower cleavage products with the full protein levels generated a ratio of CD44 cleavage/total. Vehicle: DMSO. Results are mean ± SEM; n=3. For statistical analysis mean values were compared using a 2-way ANOVA and Sidak post-hoc test; *p ≤ 0.05.

**Fig. S3.**
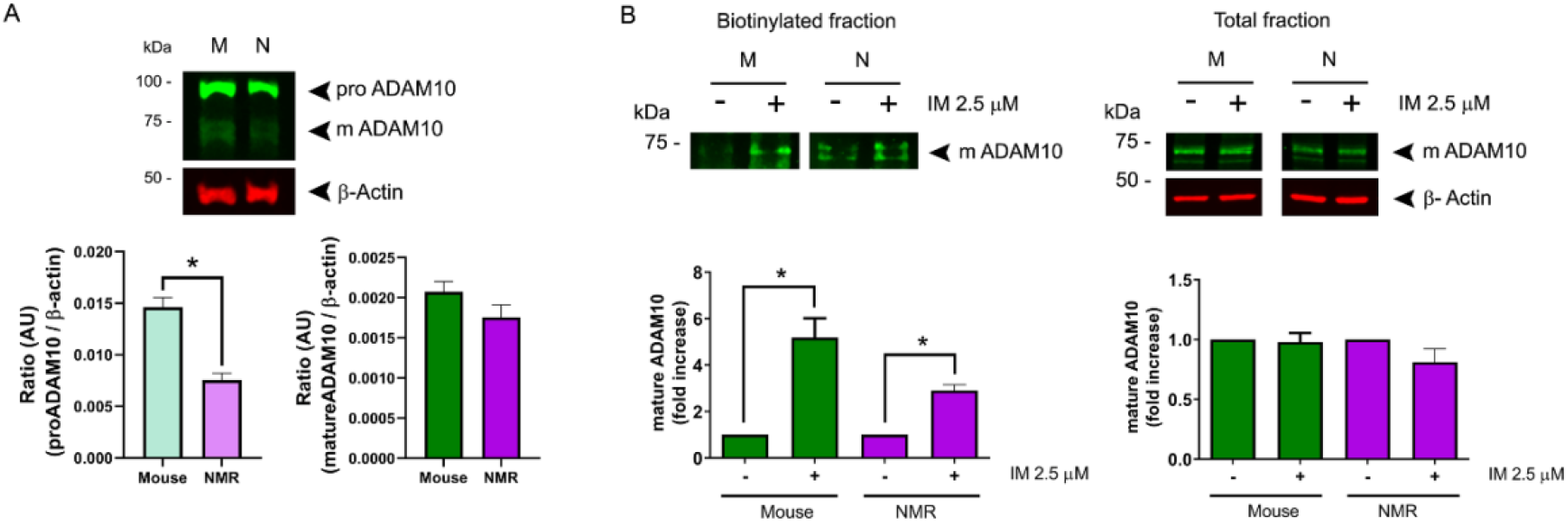
Mouse and NMR SV40-Ras skin fibroblasts express ADAM10, which undergoes IM-induced cell membrane translocation. **(A)** Immunoblot of ADAM10, where pro- and mature forms were observed in mouse and NMR SV40-Ras skin fibroblasts. β-actin was used as loading control. Results are mean ± SEM; n=3. For statistical analysis, mean values were compared using an unpaired t test; *p ≤ 0.05. **(B)** Immunoblot of mature ADAM10, comparing the biotinylated protein from the cell surface membrane fraction and total protein fraction in mouse and NMR SV40-Ras skin fibroblasts treated with ionomycin (IM, 2.5 μM). β-actin was used as loading control in the total protein fraction. Results are mean ± SEM; n=3. For statistical analysis, mean values were compared using an unpaired t test; *p ≤ 0.05.

### NMR SV40-Ras skin fibroblasts have more phosphatidylserine in the outer leaflet of the cell membrane

To try to elucidate why IM fails to induce ADAM10 shedding activity in NPSF, but does in NMR SV40-Ras skin fibroblasts, the localisation of the phospholipid phosphatidylserine (PS) in the outer leaflet of the cell membrane was evaluated. This is because surface exposure of PS on the cell membrane has been proposed to regulate ADAM10 shedding activity (*43*). Using the RealTime-Glo™ Annexin V luminescence assay, the binding of annexin V with PS in the outer leaflet of the cell membrane was measured, providing a readout in relative luminescence units (RLU). Simultaneous staining of the nucleus with the NucRed™ Live 647 ReadyProbes^®^ reagent provided a readout of relative fluorescence units (RFU), indicative of the number of cells. The ratio of both parameters was determined in MPSF, NPSF, mouse and NMR SV40-Ras skin fibroblasts (Fig. 4A). No differences were found in the PS exposure to the outer leaflet in MPSF versus mouse SV40-Ras cells, however, a higher level of PS was found in NMR SV40-Ras cells compared to NPSF (Fig. 4B). This difference in PS levels could explain why IM-induced ADAM10 shedding was observed in NMR SV40-Ras, but not NPSF.

**Fig. 4.**
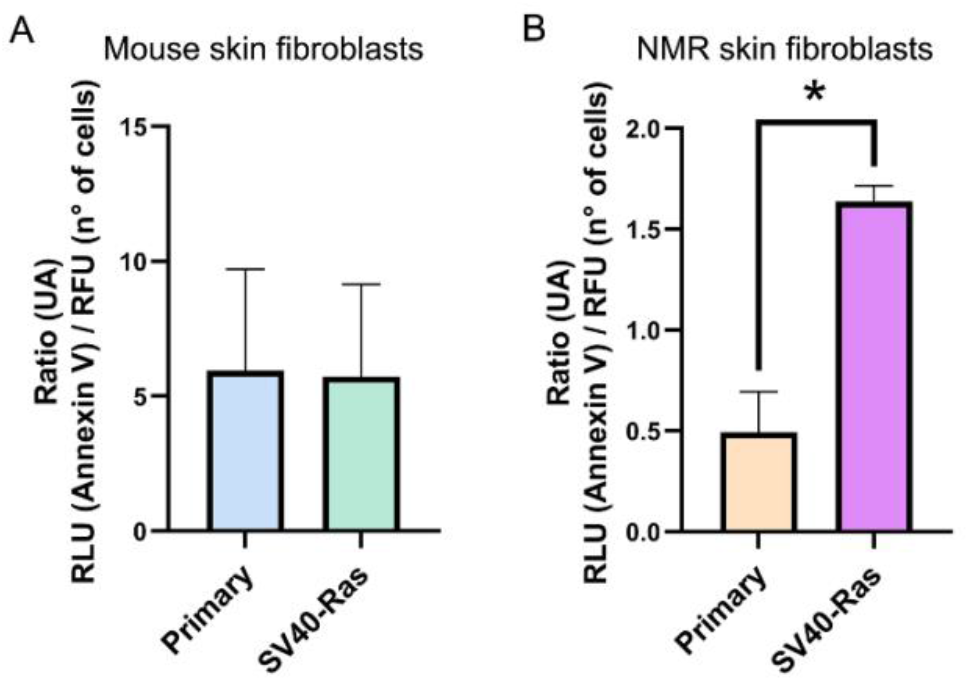
NMR SV40-Ras skin fibroblasts have more phosphatidylserine in the outer leaflet of their cell membranes than NPSF. PS in the outer leaflet of the cell membrane was evaluated through the ratio of Annexin V (RLU) and number of cells (RFU) in **(A)** mouse and **(B)** NMR, primary and SV40-Ras skin fibroblasts. Results are mean ± SEM; n=3. For statistical analysis, mean values were compared using an unpaired t test; *p ≤ 0.05.

### Overexpression of ANO6-HA rescues IM-induced ADAM10 shedding activity in NMR primary skin fibroblasts

To test whether increasing PS in the outer leaflet of the cell membrane could enhance IM-induced ADAM10 shedding activity, MPSF and NPSF were transfected with either a hyperactive form of anoctamin 6 (ANO6-HA) or GFP as a control. ANO6, is a Ca^2+^-dependent phospholipid scramblase that increases PS externalization and ADAM10-dependent substrate shedding in COS7 cells (*43*). 24 h post-transfection, increased ANO6 was observed in both MPSF and NPSF as evaluated by immunoblotting (Fig. 5, A and D). Concomitant with the ANO6 increase, an increase in PS levels in the outer leaflet of the cell membrane was observed in both MPSF and NPSF (Fig. 5, B and E). Next, MPSF and NPSF were cotransfected with AP-tagged BTC and ANO6-HA (or GFP as control) for 24 h before evaluating ADAM10 shedding activity. As expected, treatment for 1 h with 0.5 μM IM induced an increase in BTC shedding in both control and ANO6-HA MPSF, which was inhibited by batimastat (BB, 10 μM) and GI254023X (GI, 1 μM); IM-induced BTC shedding was higher in cells overexpressing ANO6-HA (Fig. 5C). Furthermore, as shown previously (Fig. 2D), in NPSF cotransfected with AP-tagged BTC and GFP, IM 0.5 μM did not induce BTC shedding, however, in cells cotransfected with AP-tagged BTC and ANO6-HA, IM induced BTC shedding, which was inhibited by batimastat (BB, 10 μM) and GI254023X (GI, 1 μM) (Fig. 5F). Due to NPSF not surviving at 37°C / 20% O_2_ (*51*), we considered if the difference in culturing conditions could explain the absence of IM-induced ADAM10 shedding activity in NPSF. Therefore, we performed the same experiments with MPSF cultured in the same conditions as NPSF. An increase of PS in the outer leaflet of the cell membrane was found in 32°C / 3% O_2_ MPSF transfected with the ANO6-HA compared with control (Fig. S4A). Moreover, unlike with NPSF, when cultured at 32°C / 3% O_2_, MPSF cotransfected with AP-tagged BTC and GFP showed increased BTC shedding following IM treatment, which, as with MPSF at 37°C, was greater in cells overexpressing ANO6-HA (Fig. S4B). Therefore, the culturing conditions alone cannot explain the difference in responses to IM between NPSF and MPSF. To determine if the higher levels of PS in the outer leaflet of the cell membrane found in NMR SV40-Ras skin fibroblasts compared to NPSF might be due to higher expression of the scramblase ANO6, protein levels of ANO6 were evaluated by immunoblot in mouse and NMR primary and SV40-Ras skin fibroblasts, however no difference was found (Fig. S5). These results demonstrate that NMR ADAM10 is a functional metalloprotease and that the lower level of PS in the outer leaflet of the cell membrane, rather than culturing conditions, is likely responsible for the lack of IM-induced ADAM10 shedding activity in NPSF.

**Fig. 5.**
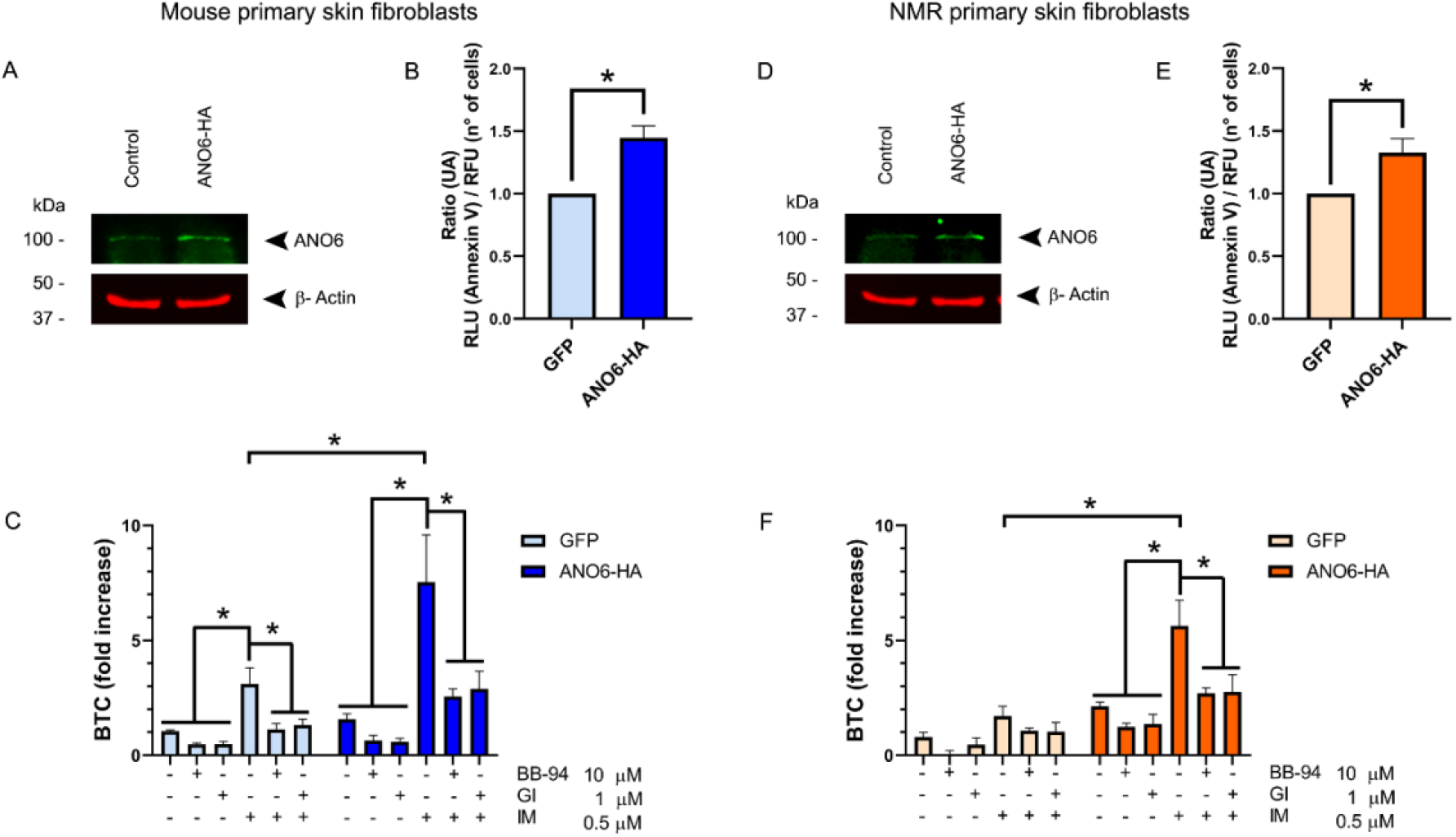
Overexpression of ANO6-HA rescues IM-induced ADAM10 shedding activity in NMR primary skin fibroblasts. Immunoblot of ANO6 in untransfected and ANO6 hyperactive (ANO6-HA) transfected **(A)** MPSF and **(D)** NPSF. β-actin was used as loading control. PS in the outer leaflet of the cell membrane was evaluated through the ratio between the Annexin V (RLU) and number of cells (RFU) in cells transfected with ANO6-HA or GFP, as a control, in **(B)** MPSF and **(E)** NPSF. Results are mean ± SEM; n=4. For statistical analysis, mean values were compared using an unpaired t test; *p ≤ 0.05. **(C)** MPSF and **(F)** NPSF were cotransfected with the AP-tagged ADAM10 substrate BTC and ANO6-HA or GFP. BTC shedding was evaluated after cells were treated for 30 min with a general inhibitor of metalloproteases (batimastat, BB, 10 μM) or the ADAM10 inhibitor (GI254023X, GI, 1 μM), and then cells were stimulated for 1 h with ionomycin (IM, 0.5 μM). Vehicle: DMSO. Results are mean ± SEM; n=4 for MPSF and n=3 for NPSF. For statistical analysis, mean values were compared using an ANOVA and Tukey’s post-hoc test; *p ≤ 0.05.

**Fig. S4.**
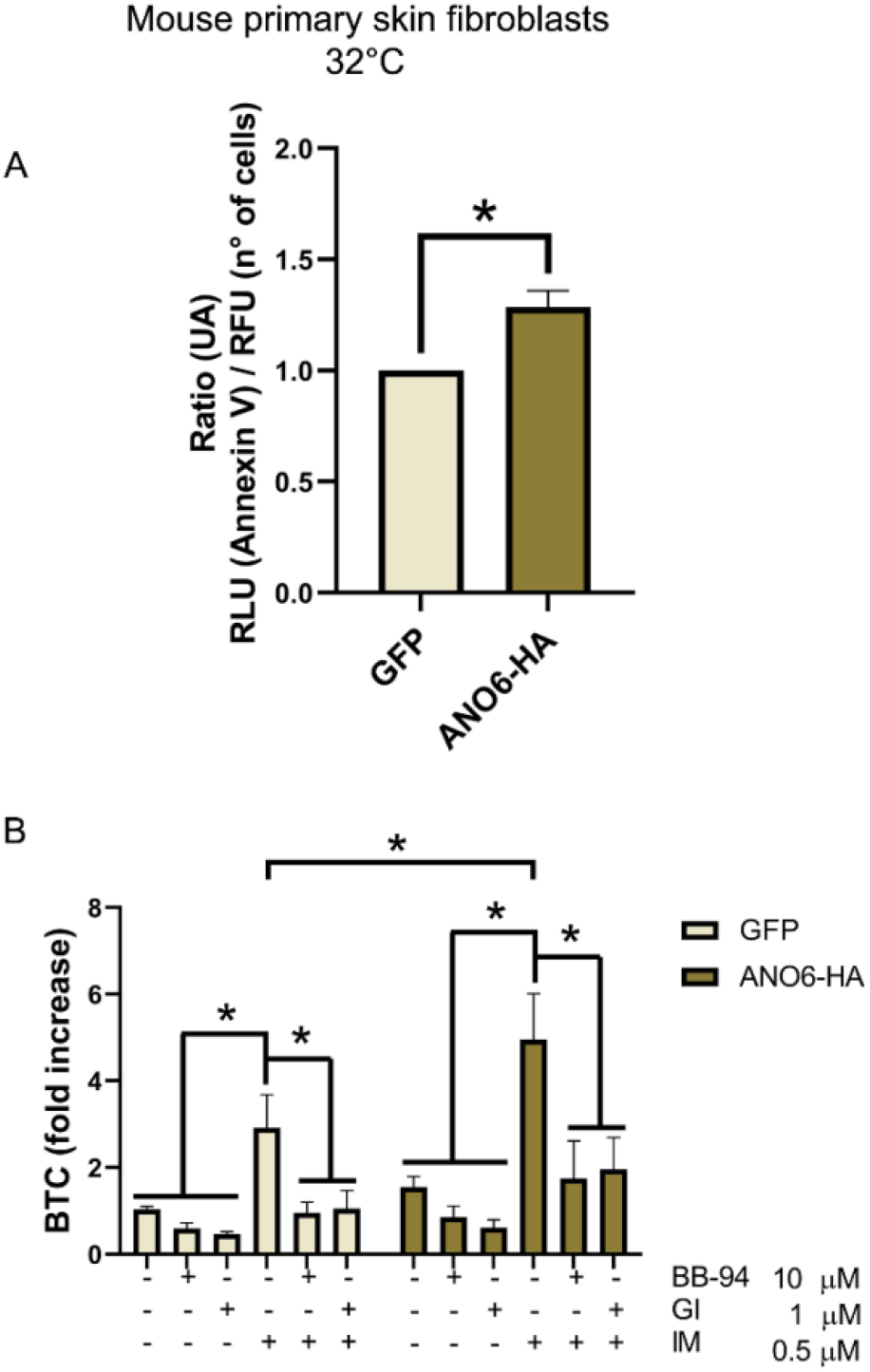
Overexpression of ANO6-HA enhances IM-induced ADAM10 shedding activity in MPSF incubated at 32°C and hypoxic conditions. **(A)** PS in the outer leaflet of the cell membrane was evaluated through the ratio between the Annexin V (RLU) and number of cells (RFU) in cells transfected with ANO6-HA or GFP as a control, in MPSF incubated at 32°C under hypoxic conditions. Results are mean ± SEM; n=3. For statistical analysis, mean values were compared using an unpaired t test; *p ≤ 0.05. **(B)** MPSF were transfected with the AP-tagged ADAM10 substrate BTC or GFP. BTC shedding was evaluated by treating cells for 30 min with a general inhibitor of metalloproteases (batimastat, BB, 10 μM) or an ADAM10 inhibitor (GI254023X, GI, 1 μM), and then cells were stimulated for 1 h with ionomycin (IM, 0.5 μM). Vehicle: DMSO. Results are mean ± SEM; n=4. For statistical analysis, mean values were compared using an ANOVA and Tukey’s post-hoc test; *p ≤ 0.05.

**Fig. S5.**
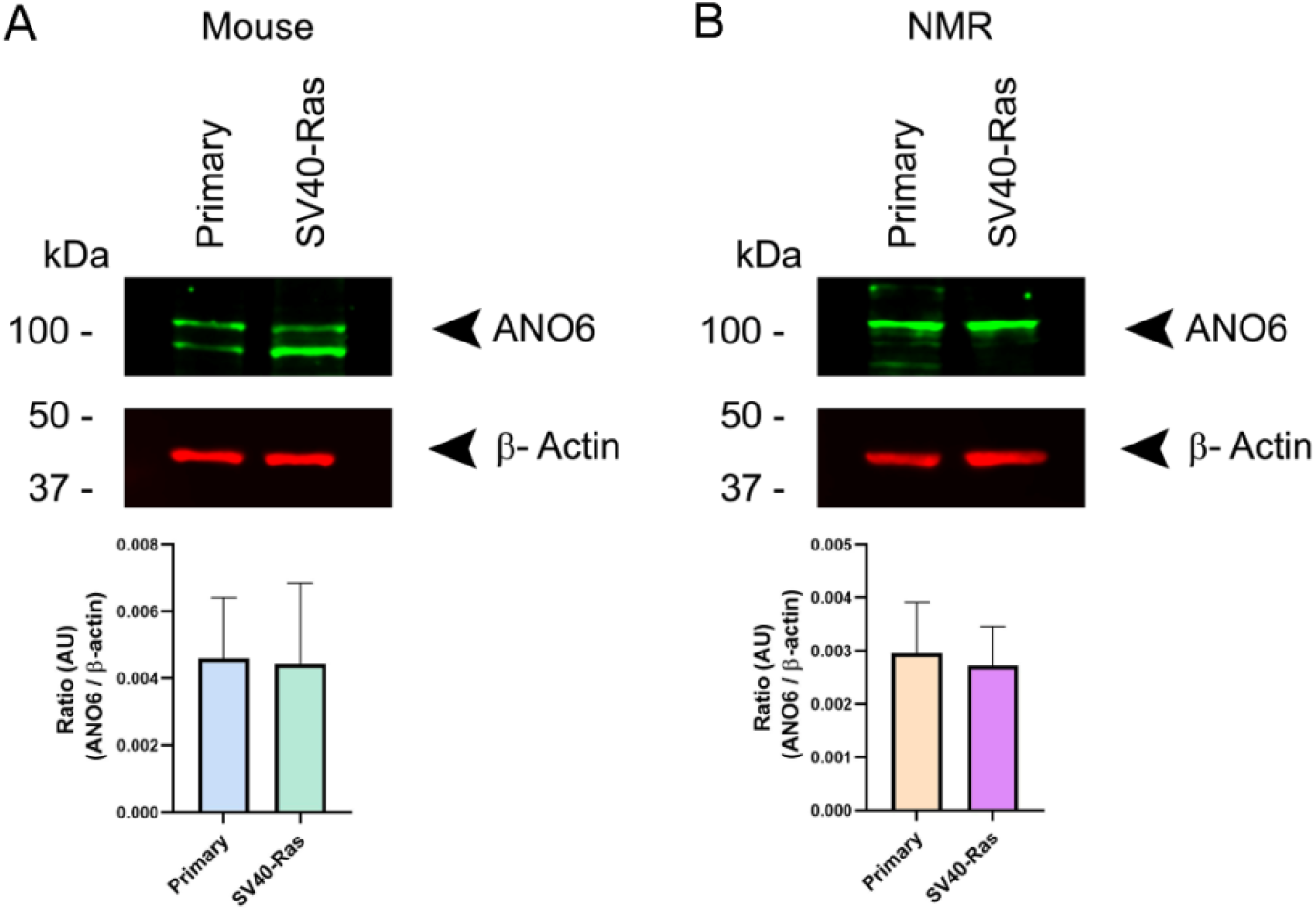
ANO6 protein levels in mouse and NMR fibroblasts. Immunoblot of ANO6, where was observed in **(A)** MPSF and Mouse SV40-Ras skin fibroblasts, and **(B)** NPSF and NMR SV40-Ras. β-actin was used as loading control. Results are mean ± SEM; n=3 for mouse and n=6 for NMR. For statistical analysis, mean values were compared using an unpaired t test.

### ADAM10 is involved in the migration of NPSF enhanced by the overexpression of ANO6-HA

Migration of tumour cells is necessary for tumour-cell invasion and metastasis. CD44 signalling has been shown to be involved in cell migration in different cells models, including glioma cells (*52*) and several breast cancer cell lines (*53*). In addition, extracellular cleavage of CD44 promotes migration in different models such as, the pancreatic tumour cell line, MIA PaCa-2 (*54*), human lung adenocarcinoma A549 cells (*29*), and certain human breast carcinoma cell lines (*30*). Interestingly, CD44 can induce its own cleavage by ADAM10 through activating the Rac pathway, as demonstrated in migration studies using glioblastoma and lung adenocarcinoma cell lines (*29*, *55*). Therefore, it was next evaluated if overexpression of ANO6-HA in NPSF could enhance cell migration. NPSF were seeded and transfected in a dish with a silicone insert. After 24 h the insert was removed to create a gap or “wound”, and images were taken regularly to measure the percentage of wound closure. NPSF displayed an increased percentage wound closure when transfected with ANO6-HA compared to GFP transfected cells (Fig. 6). The ANO6-HA induced increase in wound closure was prevented when NPSF were incubated with the ADAM10 inhibitor GI254023X (GI, 1 μM) (Fig. 6 dotted line), indicating that ADAM10 is involved in the migration enhanced by the ANO6-HA overexpression. The same trend was observed in MPSF, where the ADAM10 inhibitor GI254023X (GI, 1 μM) reduced the increase in wound closure induced by ANO6-HA when cells were cultured at 37°C and 32°C (Fig. S6).

**Fig. 6.**
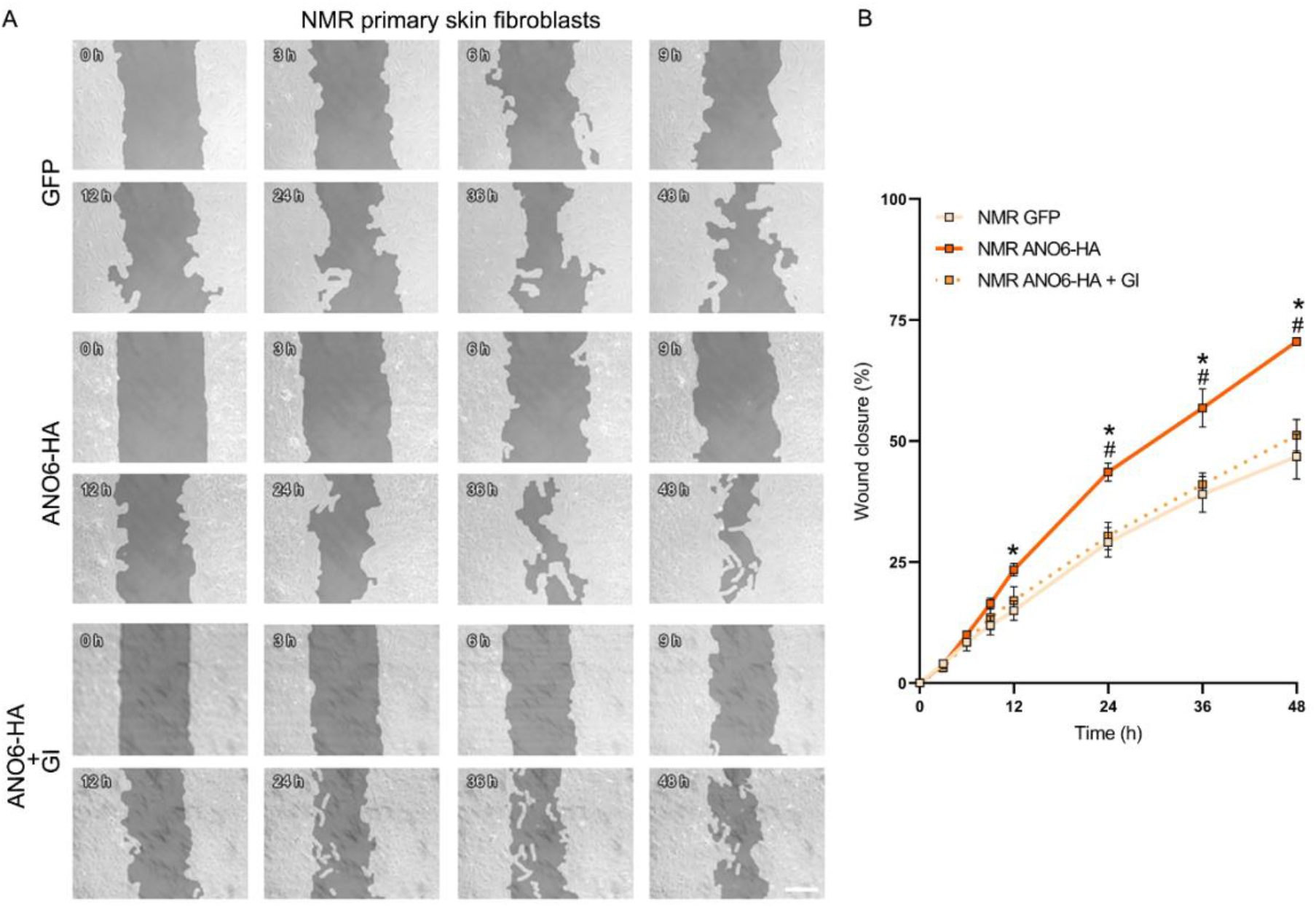
ADAM10 participates in the enhanced migration of NPSF induced by ANO6-HA overexpression. **(A)** NPSF were transfected with ANO6-HA or GFP. 24 h later the silicone insert was removed and ADAM10 inhibitor, GI254023X (GI, 1 μM) or DMSO (vehicle) was added. Images were taken at 0, 3, 6, 9,12, 24, 36, and 48 h after removal of the silicone insert for gap creation. Scale bar: 200 μm. (B) Quantification of the wound closure over the time is showed. Results are mean ± SEM; n=3. For statistical analysis, mean values were compared using a 2-way ANOVA and Tukey’s post-hoc test; *p ≤ 0.05 between GFP and ANO6-HA, #p ≤ 0.05 between ANO6-HA and ANO6-HA + GI.

**Fig. S6.**
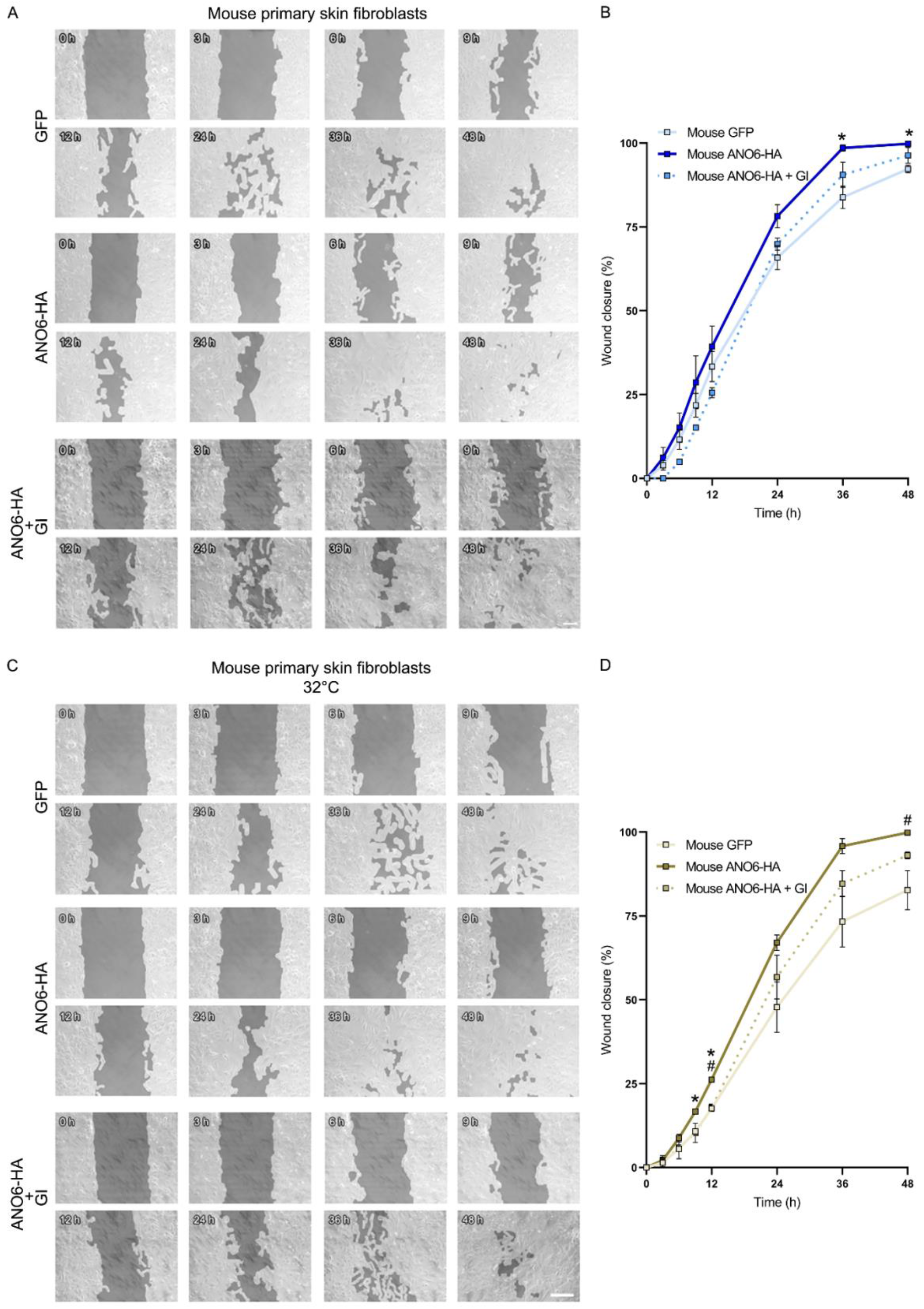
Overexpression of ANO6-HA enhances migration in mouse primary skin fibroblasts. MPSF were transfected with ANO6-HA or GFP and cultured at **(A)**37°C / 20% O_2_ or **(C)**32°C 3% O_2_. 24 h later the silicone insert was removed and ADAM10 inhibitor, GI254023X (GI, 1 μM) or DMSO (vehicle) was added. Images were taken at 0, 3, 6, 9,12, 24, 36, and 48 h after removed the silicone insert for gap creation. Scale bar: 200 μm. **(B and C)** Quantification of the wound closure over the time is showed. Results are mean ± SEM; n=4 for mouse 37°C / 20% O2 and n=3 for mouse at 32°C / 3% O2. For statistical analysis, mean values were compared using a 2-way ANOVA and Tukey’s post-hoc test; *p ≤ 0.05 between GFP and ANO6-HA, #p ≤ 0.05 between ANO6-HA and ANO6-HA + GI.

## DISCUSSION

NMRs are cancer resistant, but the mechanisms underpinning this are still not fully understood (*1*, *2*). Demonstration that NMR cells are susceptible to oncogenic transformation (*5*) suggests that other mechanisms likely explain why oncogenic events rarely result in cancer, for example a low somatic mutation rate (*6*) or altered immune response (*7*). A key part of cancer development is the ability of cells to proliferate and migrate, which involves navigating the extracellular matrix. CD44 is a receptor for several extracellular matrix components (*8*, *56*, *57*),and is considered a marker of cancer (*15*–*18*, *58*–*60*). CD44 can be cleaved in the ectodomain by metalloproteases (*20*–*22*), resulting in a sequential proteolytic cleavage of the intracellular domain by γ-secretase, generating release of the CD44 intracellular domain fragment (CD44ICD) (*61*), which translocates to the nucleus, regulating transcriptional activity (*48*). In addition, CD44ICD regulates the transcription of proteins like MMP-9 (*62*), IFN-γ (*63*) and CD44 itself (*48*).

Patients with cancer have higher CD44 cleavage levels (*25*, *26*), and ADAM10 is one metalloprotease that cleaves extracellular CD44 (*22*, *32*, *33*, *64*). Our data show that IM, which is well characterised as being able to induce ADAM10 shedding activity (*41*, *42*), only does so in transformed NMR SV40-Ras skin fibroblasts, but not in NPSF. Comparing MPSF and NPSF, we observed that IM only induces ADAM10-dependent CD44 cleavage in MPSF.

ADAM10 is of particular interest in cancer, being highly expressed, up-regulated and involved in patients with glioblastoma (*67*), lung cancer (*68*), and pancreatic cancer (*69*) to name but a few; ADAM10 overexpression also being associated with a worse prognosis for pancreatic and lung cancers (*70*). We therefore evaluated expression and shedding activity of ADAM10 to understand if the absence of IM-induced CD44 cleavage observed in NPSF was simply due to lack of ADAM10 expression. Comparing ADAM10 protein expression in MPSF and NPSF, higher levels of pro-ADAM10, but not mature ADAM10, were found in NPSF. The cleavage site of ADAM substrates is located close to the cell surface (*71*), and we found that IM induced ADAM10 translocation in both MPSF and NPSF, as well as in the transformed mouse and NMR SV40-Ras skin fibroblasts. Thus, a lack of IM-induced ADAM10-dependent CD44 cleavage is not due to lack of ADAM10 expression or aberrant plasma membrane translocation. Measuring ADAM10-dependent BTC shedding, as another approach to evaluate ADAM10 activity, we found that IM induces BTC shedding in MPSF, mouse SV40-Ras skin fibroblasts, and in NMR SV40-Ras skin fibroblasts, but not in the NPSF, results consistent with the absence of IM-induced CD44 cleavage in NPSF.

Cell membrane lipid composition regulates the shedding activity of ADAM10, such that low cholesterol levels in the cell membrane increase ADAM10 shedding activity, whereas cholesterol enrichment does not (*72*). Interestingly, the NMR brain has higher cholesterol levels than the mouse brain (*73*), which may help to explain why under basal conditions there are lower protein levels of CD44 cleavage products and also an absence of IM-induced ADAM10-dependent CD44 cleavage in NPSF. A further factor regulating ADAM10 shedding activity is PS exposure in the outer leaflet of the cell membrane. The cationic amino acid residues in the stalk region of ADAM10 are thought to interact with the negatively charged PS head group to trigger ADAM10 sheddase activity (*43*). Using mass spectrometry, no difference in the concentration of PS from NMR and mouse total brain lipid extract was detected (*73*). However, here we found approximately 3-fold higher levels of PS exposure on the cell surface of transformed NMR SV40-Ras skin fibroblasts compared to NPSF, but no changes in the PS externalization in mouse SV40-Ras compared with the MPSF. By comparison, PS exposure on the outer leaflet of the cell membrane in NPSF is only about 10% compared to the level in MPSF. This lower level of PS externalization found in NPSF could explain the lower basal protein levels of CD44 cleavage products and the absence of IM-induced CD44 cleavage and ADAM10 shedding activity in NPSF. Consistent with this, when we overexpressed a hyperactive form of ANO6, a scramblase that regulates the PS exposure in the outer leaflet of the cell membrane (*43*), we observed an increase in the PS exposure in NPSF, and as expected, IM-induced ADAM10 shedding activity was “rescued” in these cells as evaluated by the shedding of BTC. Although a higher level of PS exposure at the cell membrane was found in NMR SV40-Ras cells compared to NPSF, no difference was observed in the protein levels of ANO6, one of the scramblases that is responsible of the exposure of PS to the outer leaflet of the cell membrane (*74*). However, the abundance of ANO6 is not the only factor to consider and further analysis is required to determine if perhaps processes regulating ANO6 activation are different between cell types. Furthermore, PS exposure is not only dependent on ANO6, other scramblases, such as PLSCR (*75*) and XKR family members also participating in this process (*76*). Moreover, there are also a plethora of flippases and floppases that regulate phospholipid translocation, and thus might also contribute to the raised PS levels observed in NMR SV40-Ras cells compared to NPSF (*77*).

It has been shown that CD44 cleavage promotes tumour cell migration (*24*, *29*–*31*) and that ADAM10-dependent CD44 cleavage promotes migration of pituitary adenoma (*64*), glioblastoma (*55*), and melanoma cells (*32*). In line with these findings, in a cell migration assay, we observed that the percentage wound closure in NPSF was lower than that of MPSF, but that higher closure occurred following overexpression of ANO6-HA, which was prevented by the ADAM10 inhibitor, GI254023X, thus suggesting that ADAM10 shedding activity might contribute to the NPSF migration.

Taking all these data together, our findings suggest that the lower level of PS in the outer membrane of NPSF prevents ADAM10 shedding activity in response to IM. Future work will be focused on understanding the regulation of PS localization in NPSF, including the roles of scramblases, flippases and floppases, as well as studying the regulation and localization of Ca^2+^ in the NPSF compared to MPSF. To fully establish whether diminished ADAM10 activity in MPSF contributes to their cancer resistance, *in vivo* studies would be the next appropriate step.

## Acknowledgments

The plasmid for alkaline phosphatase (AP)-tagged BTC expression was a kind gift from Dr Carl P. Blobel (Hospital for Special Surgery, New York, USA). The plasmid for the hyperactive form of ANO6 (ANO6-HA) was a kind gift from Dr Karina Reiß (Department of Dermatology, University of Kiel, 24105 Kiel, Germany). For the purpose of open access, the author has applied a Creative Commons Attribution (CC BY) licence to any Author Accepted Manuscript version arising from this submission

## Author contributions

**Paulina Urriola-Munoz:** conceptualization, formal analysis, investigation, methodology, project administration, validation, visualization, writing – original draft and writing – review & editing. **Luke A. Pattison:** investigation. **Ewan St. John. Smith:** conceptualization, funding acquisition, methodology, supervision and writing – review & editing.

## Data and materials availability

The data that support the findings of this study are available from the corresponding author upon reasonable request.

## REFERENCES

1. R. Buffenstein et al., The naked truth: a comprehensive clarification and classification of current ‘myths’ in naked mole-rat biology. Biol Rev Camb Philos Soc 97, 115–140 (2022).

2. F. Hadi, E. S. J. Smith, W. T. Khaled, Naked Mole-Rats: Resistant to Developing Cancer or Good at Avoiding It? Adv Exp Med Biol 1319, 341–352 (2021).

3. Y. H. Edrey, M. Hanes, M. Pinto, J. Mele, R. Buffenstein, Successful aging and sustained good health in the naked mole rat: a long-lived mammalian model for biogerontology and biomedical research. ILAR J 52, 41–53 (2011).

4. R. Buffenstein, The naked mole-rat: a new long-living model for human aging research. J Gerontol A Biol Sci Med Sci 60, 1369–1377 (2005).

5. F. Hadi et al., Transformation of naked mole-rat cells. Nature 583, E1–E7 (2020).

6. A. Cagan et al., Somatic mutation rates scale with lifespan across mammals. Nature 604, 517–524 (2022).

7. K. Oka et al., Resistance to chemical carcinogenesis induction via a dampened inflammatory response in naked mole-rats. Commun Biol 5, 287 (2022).

8. B. Radotra, D. McCormick, A. Crockard, CD44 plays a role in adhesive interactions between glioma cells and extracellular matrix components. Neuropathol Appl Neurobiol 20, 399–405 (1994).

9. S. Jalkanen, M. Jalkanen, Lymphocyte CD44 binds the COOH-terminal heparin-binding domain of fibronectin. J Cell Biol 116, 817–825 (1992).

10. K. J. Wolf et al., A mode of cell adhesion and migration facilitated by CD44-dependent microtentacles. Proc Natl Acad Sci U S A 117, 11432–11443 (2020).

11. N. Ludwig et al., CD44(+) tumor cells promote early angiogenesis in head and neck squamous cell carcinoma. Cancer Lett 467, 85–95 (2019).

12. T. Yae et al., Alternative splicing of CD44 mRNA by ESRP1 enhances lung colonization of metastatic cancer cell. Nat Commun 3, 883 (2012).

13. M. Zoller, CD44: physiological expression of distinct isoforms as evidence for organ-specific metastasis formation. J Mol Med (Berl) 73, 425–438 (1995).

14. L. Y. Bourguignon, Hyaluronan-mediated CD44 activation of RhoGTPase signaling and cytoskeleton function promotes tumor progression. Semin Cancer Biol 18, 251–259 (2008).

15. M. Hassn Mesrati, S. E. Syafruddin, M. A. Mohtar, A. Syahir, CD44: A Multifunctional Mediator of Cancer Progression. Biomolecules 11, (2021).

16. S. Takaishi et al., Identification of gastric cancer stem cells using the cell surface marker CD44. Stem Cells 27, 1006–1020 (2009).

17. E. L. Leung et al., Non-small cell lung cancer cells expressing CD44 are enriched for stem cell-like properties. PLoS One 5, e14062 (2010).

18. B. L. Lokeshwar, V. B. Lokeshwar, N. L. Block, Expression of CD44 in prostate cancer cells: association with cell proliferation and invasive potential. Anticancer Res 15, 1191–1198 (1995).

19. J. Lesley, R. Hyman, CD44 structure and function. Front Biosci 3, d616–630 (1998).

20. O. Nagano, H. Saya, Mechanism and biological significance of CD44 cleavage. Cancer Sci 95, 930–935 (2004).

21. H. Nakamura et al., Constitutive and induced CD44 shedding by ADAM-like proteases and membrane-type 1 matrix metalloproteinase. Cancer Res 64, 876–882 (2004).

22. O. Nagano et al., Cell-matrix interaction via CD44 is independently regulated by different metalloproteinases activated in response to extracellular Ca(2+) influx and PKC activation. J Cell Biol 165, 893–902 (2004).

23. I. Okamoto et al., Regulated CD44 cleavage under the control of protein kinase C, calcium influx, and the Rho family of small G proteins. J Biol Chem 274, 25525–25534 (1999).

24. Y. Kawano et al., Ras oncoprotein induces CD44 cleavage through phosphoinositide 3-OH kinase and the rho family of small G proteins. J Biol Chem 275, 29628–29635 (2000).

25. I. Okamoto et al., Proteolytic cleavage of the CD44 adhesion molecule in multiple human tumors. Am J Pathol 160, 441–447 (2002).

26. N. Yamane, S. Tsujitani, M. Makino, M. Maeta, N. Kaibara, Soluble CD44 variant 6 as a prognostic indicator in patients with colorectal cancer. Oncology 56, 232–238 (1999).

27. Y. J. Guo et al., Potential use of soluble CD44 in serum as indicator of tumor burden and metastasis in patients with gastric or colon cancer. Cancer Res 54, 422–426 (1994).

28. D. Masson et al., Soluble CD44: quantification and molecular repartition in plasma of patients with colorectal cancer. Br J Cancer 80, 1995–2000 (1999).

29. C. Kolliopoulos, A. Chatzopoulos, S. S. Skandalis, C. H. Heldin, P. Heldin, TRAF4/6 Is Needed for CD44 Cleavage and Migration via RAC1 Activation. Cancers (Basel) 13, (2021).

30. C. I. Kung et al., Enhanced membrane-type 1 matrix metalloproteinase expression by hyaluronan oligosaccharides in breast cancer cells facilitates CD44 cleavage and tumor cell migration. Oncol Rep 28, 1808–1814 (2012).

31. K. N. Sugahara et al., Hyaluronan oligosaccharides induce CD44 cleavage and promote cell migration in CD44-expressing tumor cells. J Biol Chem 278, 32259–32265 (2003).

32. U. Anderegg et al., ADAM10 is the constitutive functional sheddase of CD44 in human melanoma cells. J Invest Dermatol 129, 1471–1482 (2009).

33. T. Murai, T. Miyauchi, T. Yanagida, Y. Sako, Epidermal growth factor-regulated activation of Rac GTPase enhances CD44 cleavage by metalloproteinase disintegrin ADAM10. Biochem J 395, 65–71 (2006).

34. K. Reiss, P. Saftig, The “a disintegrin and metalloprotease” (ADAM) family of sheddases: physiological and cellular functions. Semin Cell Dev Biol 20, 126–137 (2009).

35. U. Sahin et al., Distinct roles for ADAM10 and ADAM17 in ectodomain shedding of six EGFR ligands. J Cell Biol 164, 769–779 (2004).

36. D. Hartmann et al., The disintegrin/metalloprotease ADAM 10 is essential for Notch signalling but not for alpha-secretase activity in fibroblasts. Hum Mol Genet 11, 2615–2624 (2002).

37. K. Reiss et al., ADAM10 cleavage of N-cadherin and regulation of cell-cell adhesion and beta-catenin nuclear signalling. EMBO J 24, 742–752 (2005).

38. S. B. Lee et al., ADAM10 is upregulated in melanoma metastasis compared with primary melanoma. J Invest Dermatol 130, 763–773 (2010).

39. G. Murphy, The ADAMs: signalling scissors in the tumour microenvironment. Nat Rev Cancer 8, 929–941 (2008).

40. E. Marcello, B. Borroni, S. Pelucchi, F. Gardoni, M. Di Luca, ADAM10 as a therapeutic target for brain diseases: from developmental disorders to Alzheimer’s disease. Expert Opin Ther Targets 21, 1017–1026 (2017).

41. T. Maretzky et al., ADAM10 mediates E-cadherin shedding and regulates epithelial cell-cell adhesion, migration, and beta-catenin translocation. Proc Natl Acad Sci U S A 102, 9182–9187 (2005).

42. K. Reiss, S. Bhakdi, Pore-forming bacterial toxins and antimicrobial peptides as modulators of ADAM function. Med Microbiol Immunol 201, 419–426 (2012).

43. F. Bleibaum et al., ADAM10 sheddase activation is controlled by cell membrane asymmetry. J Mol Cell Biol 11, 979–993 (2019).

44. E. Johansson et al., CD44 Interacts with HIF-2alpha to Modulate the Hypoxic Phenotype of Perinecrotic and Perivascular Glioma Cells. Cell Rep 20, 1641–1653 (2017).

45. K. Horiuchi et al., Substrate selectivity of epidermal growth factor-receptor ligand sheddases and their regulation by phorbol esters and calcium influx. Mol Biol Cell 18, 176–188 (2007).

46. T. Geback, M. M. Schulz, P. Koumoutsakos, M. Detmar, TScratch: a novel and simple software tool for automated analysis of monolayer wound healing assays. Biotechniques 46, 265–274 (2009).

47. I. Okamoto et al., CD44 cleavage induced by a membrane-associated metalloprotease plays a critical role in tumor cell migration. Oncogene 18, 1435–1446 (1999).

48. I. Okamoto et al., Proteolytic release of CD44 intracellular domain and its role in the CD44 signaling pathway. J Cell Biol 155, 755–762 (2001).

49. A. Ludwig et al., Metalloproteinase inhibitors for the disintegrin-like metalloproteinases ADAM10 and ADAM17 that differentially block constitutive and phorbol ester-inducible shedding of cell surface molecules. Comb Chem High Throughput Screen 8, 161–171 (2005).

50. P. Urriola-Munoz et al., The xenoestrogens biphenol-A and nonylphenol differentially regulate metalloprotease-mediated shedding of EGFR ligands. J Cell Physiol 233, 2247–2256 (2018).

51. D. Omerbasic et al., Hypofunctional TrkA Accounts for the Absence of Pain Sensitization in the African Naked Mole-Rat. Cell Rep 17, 748–758 (2016).

52. S. Koochekpour, G. J. Pilkington, A. Merzak, Hyaluronic acid/CD44H interaction induces cell detachment and stimulates migration and invasion of human glioma cells in vitro. Int J Cancer 63, 450–454 (1995).

53. K. Zen et al., CD44v4 is a major E-selectin ligand that mediates breast cancer cell transendothelial migration. PLoS One 3, e1826 (2008).

54. M. Kajita et al., Membrane-type 1 matrix metalloproteinase cleaves CD44 and promotes cell migration. J Cell Biol 153, 893–904 (2001).

55. T. Murai et al., Engagement of CD44 promotes Rac activation and CD44 cleavage during tumor cell migration. J Biol Chem 279, 4541–4550 (2004).

56. K. L. Chen, D. Li, T. X. Lu, S. W. Chang, Structural Characterization of the CD44 Stem Region for Standard and Cancer-Associated Isoforms. Int J Mol Sci 21, (2020).

57. S. Banerji et al., Structures of the Cd44-hyaluronan complex provide insight into a fundamental carbohydrate-protein interaction. Nat Struct Mol Biol 14, 234–239 (2007).

58. E. Olsson et al., CD44 isoforms are heterogeneously expressed in breast cancer and correlate with tumor subtypes and cancer stem cell markers. BMC Cancer 11, 418 (2011).

59. M. D. Corte et al., Analysis of the expression of hyaluronan in intraductal and invasive carcinomas of the breast. J Cancer Res Clin Oncol 136, 745–750 (2010).

60. M. Zoller, CD44: can a cancer-initiating cell profit from an abundantly expressed molecule? Nat Rev Cancer 11, 254–267 (2011).

61. D. Murakami et al., Presenilin-dependent gamma-secretase activity mediates the intramembranous cleavage of CD44. Oncogene 22, 1511–1516 (2003).

62. K. E. Miletti-Gonzalez et al., Identification of function for CD44 intracytoplasmic domain (CD44-ICD): modulation of matrix metalloproteinase 9 (MMP-9) transcription via novel promoter response element. J Biol Chem 287, 18995–19007 (2012).

63. K. Schultz et al., Gamma secretase dependent release of the CD44 cytoplasmic tail upregulates IFI16 in cd44-/-tumor cells, MEFs and macrophages. PLoS One 13, e0207358 (2018).

64. Y. Pan et al., ADAM10 promotes pituitary adenoma cell migration by regulating cleavage of CD44 and L1. J Mol Endocrinol 49, 21–33 (2012).

65. A. Seluanov et al., Hypersensitivity to contact inhibition provides a clue to cancer resistance of naked mole-rat. Proc Natl Acad Sci U S A 106, 19352–19357 (2009).

66. X. Tian et al., High-molecular-mass hyaluronan mediates the cancer resistance of the naked mole rat. Nature 499, 346–349 (2013).

67. E. J. Siney et al., Metalloproteinases ADAM10 and ADAM17 Mediate Migration and Differentiation in Glioblastoma Sphere-Forming Cells. Mol Neurobiol 54, 3893–3905 (2017).

68. J. Guo et al., ADAM10 overexpression in human non-small cell lung cancer correlates with cell migration and invasion through the activation of the Notch1 signaling pathway. Oncol Rep 28, 1709–1718 (2012).

69. M. M. Gaida et al., Expression of A disintegrin and metalloprotease 10 in pancreatic carcinoma. Int J Mol Med 26, 281–288 (2010).

70. M. Trerotola et al., Trop-2 cleavage by ADAM10 is an activator switch for cancer growth and metastasis. Neoplasia 23, 415–428 (2021).

71. K. Horiuchi, A brief history of tumor necrosis factor alpha--converting enzyme: an overview of ectodomain shedding. Keio J Med 62, 29–36 (2013).

72. V. Matthews et al., Cellular cholesterol depletion triggers shedding of the human interleukin-6 receptor by ADAM10 and ADAM17 (TACE). J Biol Chem 278, 38829–38839 (2003).

73. D. Frankel et al., Cholesterol-rich naked mole-rat brain lipid membranes are susceptible to amyloid beta-induced damage in vitro. Aging (Albany NY) 12, 22266–22290 (2020).

74. J. Suzuki, M. Umeda, P. J. Sims, S. Nagata, Calcium-dependent phospholipid scrambling by TMEM16F. Nature 468, 834–838 (2010).

75. Q. Zhou et al., Molecular cloning of human plasma membrane phospholipid scramblase. A protein mediating transbilayer movement of plasma membrane phospholipids. J Biol Chem 272, 18240–18244 (1997).

76. J. Suzuki, D. P. Denning, E. Imanishi, H. R. Horvitz, S. Nagata, Xk-related protein 8 and CED-8 promote phosphatidylserine exposure in apoptotic cells. Science 341, 403–406 (2013).

77. D. L. Daleke, Regulation of transbilayer plasma membrane phospholipid asymmetry. J Lipid Res 44, 233–242 (2003).

